# Charting the cellular biogeography in colitis reveals fibroblast trajectories and coordinated spatial remodeling

**DOI:** 10.1101/2023.05.08.539701

**Authors:** Paolo Cadinu, Kisha N. Sivanathan, Aditya Misra, Rosalind J. Xu, Davide Mangani, Evan Yang, Joseph M. Rone, Katherine Tooley, Yoon-Chul Kye, Lloyd Bod, Ludwig Geistlinger, Tyrone Lee, Noriaki Ono, Gang Wang, Liliana Sanmarco, Francisco J. Quintana, Ana C. Anderson, Vijay K. Kuchroo, Jeffrey R. Moffitt, Roni Nowarski

## Abstract

Gut inflammation involves contributions from immune and non-immune cells, whose interactions are shaped by the spatial organization of the healthy gut and its remodeling during inflammation. The crosstalk between fibroblasts and immune cells is an important axis in this process, but our understanding has been challenged by incomplete cell-type definition and biogeography. To address this challenge, we used MERFISH to profile the expression of 940 genes in 1.35 million cells imaged across the onset and recovery from a mouse colitis model. We identified diverse cell populations; charted their spatial organization; and revealed their polarization or recruitment in inflammation. We found a staged progression of inflammation-associated tissue neighborhoods defined, in part, by multiple inflammation-associated fibroblasts, with unique expression profiles, spatial localization, cell-cell interactions, and healthy fibroblast origins. Similar signatures in ulcerative colitis suggest conserved human processes. Broadly, we provide a framework for understanding inflammation-induced remodeling in the gut and other tissues.

## INTRODUCTION

Intestinal tissue organizes a remarkable variety of functions ranging from nutrient absorption and processing, to environmental sensation, to establishing the balance between immune tolerance and inflammation^1–3^. These functions, in turn, are executed by a diverse range of cell types whose individual and cooperative functions collectively give rise to gut homeostasis. In the context of Inflammatory Bowel Diseases (IBD), such as Crohn’s disease (CD) and ulcerative colitis (UC)—chronic disorders characterized by relapsing and remitting inflammation of the intestine—the cellular diversity of the healthy gut is remodeled. This process involves not only recruitment of new immune cell populations, but also the polarization of new inflammation-associated states of existing populations^3–8^. Thus, a better understanding of this cellular landscape in both the healthy and inflamed gut may facilitate a deeper understanding of and new therapeutic avenues for IBD.

For this reason, recent single-cell RNA sequencing (scRNAseq) studies cataloguing stromal^9–18^, epithelial^19–25^, immune^26–29^, and multiple^30–41^ cell populations in the healthy and inflamed gut have generated new insights in both mice and humans. In addition to providing transcriptome-wide expression profiles for established cell types, these studies have helped define novel and less well appreciated populations or functions, including, among many, repair-associated epithelial populations^24, 25^, goblet cell subsets^22^, and immunoregulatory functions of non-immune cells^16^. However, these studies have arguably had the biggest impact on our understanding of fibroblasts, where the molecular, functional, and phenotypic heterogeneity of these cells has only been recently appreciated^9–18, 42, 43^. While a consensus on the subclassification and naming of fibroblast populations has not yet been reached, it is clear that the small and large intestine contain multiple fibroblast populations with distinguishing gene-expression profiles, characteristic positioning in subepithelial, interstitial, pericryptal, perivascular, or follicular domains, and functionally distinct roles^9–11, 13–15, 17, 18, 43^. Moreover, it has also been recently appreciated that fibroblasts can polarize in the context of inflammation and give rise to inflammation-associated fibroblasts (IAFs)^9, 15, 17, 18, 31–33, 36, 39–41^. IAFs have been identified in CD^30, 32, 33, 36, 41^ and UC^9, 31^ biopsies and in mouse models of IBD^9, 15, 18, 39, 40^ and have proposed roles in extracellular matrix (ECM) remodeling, immune recruitment, oxidative tissue damage and inflammatory cytokine production. However, the diversity of IAF populations, their spatial location, their emergence at different stages of inflammation, and their origin in healthy fibroblast populations remains unclear.

In parallel, the intricate structural organization of the gut dictates the way in which cell populations interact, and contribute to tissue remodeling in inflammation^44, 45^. Spatial remodeling of the inflamed colonic mucosa includes crypt hyperplasia, erosion, and abscesses, culminating in tissue ulceration. In non-mucosal regions, spatial remodeling includes submucosal thickening, immune infiltration, increase in tertiary lymphoid structure abundance, and the emergence of creeping fat. Underscoring the importance of these structural changes, their quantification via histological scoring is used for IBD diagnosis and treatment prognosis^46^. Nonetheless, the cell-type diversity revealed by single-cell transcriptomics studies has not yet been fully registered within these structural changes. Illustrating the promise of such integration, a recent combination of bulk transcriptomics with histopathology in IBD biopsies implicated ulcerated tissue, and the collection of granulocytes, mononuclear phagocytes, and activated fibroblasts found within, in patient non-response to anti-TNF treatment^16^.

Image-based approaches to spatial transcriptomics^47^ provide the ability to simultaneously chart the cellular and spatial organization of tissues and promise to merge the insights provided by single-cell transcriptomics with those provided by tissue morphology. Multiplexed-error robust-fluorescence in situ hybridization (MERFISH) is one such method that can image and identify thousands of different RNA molecules within single cells within their native tissue context and, thus, define and discover cell types and states, infer cellular function, and chart tissue organization from these *in situ* measurements^48–51^. Moreover, MERFISH has been used to define cell populations or states not easily seen in scRNAseq due to its increased molecular sensitivity, higher cellular throughput, and lack of dissociation-induced transcriptional or cell-type abundance perturbations^51, 52^. Here we extend the use of MERFISH from the brain, where it has been most commonly used^51–55^, to the gastrointestinal tract in order to chart the cellular and spatial structure of the colon in a mouse IBD model. We identify a diversity of healthy and inflammation-associated cell populations, define their organization into distinct spatial domains, and chart the staged evolution of these neighborhoods across disease. Our studies reveal distinct IAF populations with unique gene expression profiles, spatial locations, disease-stage-development, potential cell-cell interactions, and origins in healthy fibroblasts. Importantly, we find evidence that both IAF heterogeneity and the spatial remodeling events we see in mouse may have analogs in human UC. Taken together, our measurements may provide a framework for the dissection of additional aspects of the gut inflammatory response, and the approaches we leverage here may prove useful in the dissection of similar responses in other tissue and disease contexts.

## RESULTS

### MERFISH maps cellular diversity in the healthy and inflamed colon

To chart the cellular and spatial remodeling that occur during the onset of and recovery from gut inflammation, we leveraged MERFISH^48–51^ to profile gene expression at different disease stages in a dextran-sodium-sulfate (DSS)-induced mouse colitis model^56–58^. We harvested the distal colon prior to DSS treatment (Day 0), early in disease (Day 3), at peak inflammation (Day 9), and after a DSS-free recovery period during which most mice recovered their pre-treatment weight (Day 21; **Figure 1A**). In parallel, we designed a MERFISH library targeting 940 genes. The first portion of the panel contained literature-derived cell-type markers included to define and spatially register cell types while the remaining panel portion included pathways important in gut homeostasis and inflammation to provide functional insight into the identified populations and their potential interactions.

**Figure 1:**
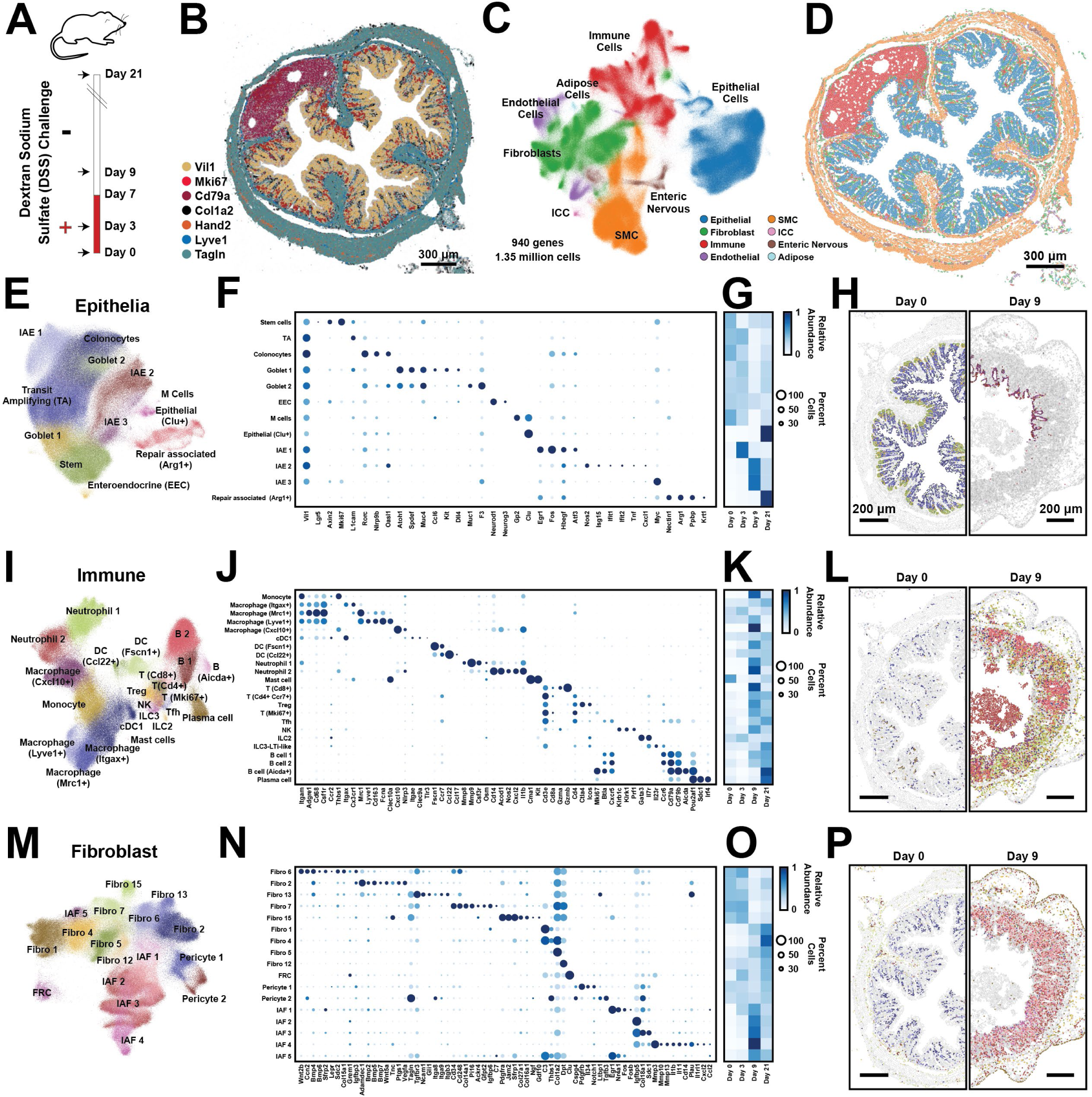
Cellular remodeling in a mouse colitis model. (A) Time-course for the administration (red) or withdrawal (white) of dextran-sodium- sulfate (DSS) in the mouse colitis model. Arrows represent the time of tissue harvest. (B) The spatial distribution of 7 of 940 RNAs measured with MERFISH in a single slice of the mouse distal colon harvested at Day 0. Dots represent individual RNA molecules and color indicates their identity. Scale bar: 300 µm. (C) Uniform Manifold Approximation and Projection (UMAP) of 1.35 million cells measured via MERFISH for mice at all harvest days. Major cell classes are indicated via color. Smooth muscle cells (SMC). Interstitial cells of Cajal (ICC). (D) The spatial distribution of major cell classes in the mouse colon slice shown in (B). Patches represent the cellular boundaries, and color indicates cell class as in (C). Scale bar: 300 µm. (E) UMAP representation of all epithelial cells with individual epithelial populations colored and labeled. (F) The expression of key genes in all identified epithelial populations. Color indicates the average expression of the listed gene and circle diameter represents the fraction of cells that express at least one copy of that RNA. Expression is normalized by the maximum value observed for each gene across all listed cell populations. (G) The average abundance of each cell population per slice across the different harvest days, normalized to the maximum abundance. (H) The spatial distribution of cells within representative portions of representative slices from Day 0 (left) and Day 9 (right). The centroid of individual cells is plotted with epithelial cells colored by their identity as in (E) and all other cells in gray. Scale bars: 200 µm (I)-(L) As in (E)-(H) but for all immune populations. (M)-(P) As in (E)-(H) but for all fibroblast populations.

To profile gene expression, the distal colon was fixed, frozen, and cryo-sectioned, and 10-µm-thick sections were stained with a novel fixed-frozen protocol developed to preserve RNA integrity. We profiled RNAs within 52 slices taken from 15 mice across these four time points and partitioned these RNAs into cells using Baysor^59^. We identified 1.7 million segmented regions and removed small cell fragments to produce 1,348,582 putative cells. On average, individual cells contained 85 ± 61 (standard deviation [STD]) RNA counts and at least one count of 57 ± 29 (STD) different genes. Supporting our measurements, we found that gene expression correlated between biological replicates and with bulk RNA sequencing (**Figures S1A**–**S1C**). Furthermore, key marker genes defined the expected anatomical gut structures, including the mucosa (comprising the epithelial layer, the lamina propria [LP] and the muscularis mucosa), the submucosa, the muscularis externa (comprising the circular and longitudinal muscle layers), the submucosal plexus and myenteric plexus, the occasional isolated lymphoid follicle, and stray serosal or mesentery fragments (**Figure 1B**).

To define cell populations, we leveraged standard single-cell analysis tools^60, 61^ and identified 72 cell clusters comprising 8 major classes—including epithelial cells; fibroblasts; immune cells; endothelial cells (EC); cells of the enteric nervous system (ENS); smooth muscle cells (SMC); adipose cells; and interstitial cells of Cajal (ICC)— which cross validate against clusters derived from published scRNAseq^9, 15, 18, 39, 40^ (**Figure 1C**-**1P**; **Figure S1D-S1K**; **Figure S2; Figure S3A-S3P**). 745 of the 940 profiled genes were measured above background in at least one cluster, confirming the informative nature of our panel. Interestingly, we noted reproducible expression gradients within many clusters with some variations cued by tissue location (**Figure S3Q-S3X**), suggesting that further meaningful cluster sub-divisions may be possible.

### DSS-induced colitis elicits broad cellular remodeling

With this atlas in place, we next explored the cellular remodeling that occurs during the onset and recovery from DSS-induced colitis by examining the clusters observed within each of the major cell classes in greater depth. Starting with the epithelial compartment in healthy mice (**Figure 1E-1H**), we identified stem cells, transit amplifying (TA) cells, and colonocytes arrayed in the expected crypt base-to-tip spatial progression. We also identified the expected secretory populations^62^ including Goblet 1 in the crypt base, Goblet 2 near the crypt tip, and enteroendocrine cells (EEC) more concentrated in the crypt base. Finally, we detected several rare epithelial populations, including M cells enriched in the follicle-associated epithelium and a recently described population of Clu+ epithelia found at the crypt bottom^24^.

Our measurements also revealed multiple epithelial populations that arose at distinct stages of inflammation, which we term inflammation-associated epithelia (IAE1, IAE2 and IAE3). IAE1 was primarily found at Day 3 and was marked by the expression of immediately early genes (IEG; e.g. *Egr1* and *Fos*; **Figure 1E-1G**) whose expression is often associated with early stages of transcriptional remodeling. Shared gene expression and location suggested that IAE1 was a polarized form of top-crypt colonocytes that up- regulated stress-inducible genes (e.g., *Atf3* and *Hbegf*) while downregulating several immune-related genes (e.g., *Nlrp9b*, *Oasl1*, and *Rorc*), suggesting that it might represent a stress-induced non-inflammatory state. IAE2 and IAE3 were more prominent at Day 9 (**Figure 1E-1G**). The expression of interferon-stimulated genes (e.g., *Isg15*, *Ifit1*, and *Ifit2*), notable inflammatory cytokines (e.g., *Tnf* and *Cxcl1*), and *Nos2*, which produces bactericidal reactive nitrogen species, suggested that IAE2 might play an inflammatory role. By contrast, the expression of high levels of *Myc* and some epithelial stem cell markers in IAE3, suggested a role in epithelial proliferation, repair, or renewal^63^. Overall, the IAE populations were largely depleted at Day 21 and were replaced with the full diversity of healthy populations with a notable increase in Clu+ epithelia relative to Day 0 (**Figure 1G**), consistent with the regenerative role of these cells^24^. In addition, Day 21 showed the emergence of a population we term Repair-associated (Arg1+). Based on their location, their lack of Vil1 expression, and their marker genes (e.g., *Arg1*, *Krt1*, and *Nectin1*; **Figure 1F**), these cells likely represent squamous epithelia shown recently to migrate to sites of epithelial damage to support repair^25^.

We also observed a wide variety of immune cells in healthy and inflamed tissue with most populations expanding substantially in disease (**Figure 1I-1L**). In the healthy colon, the most abundant immune populations were macrophages: Macrophage (Itgax+), Macrophage (Mrc1+), and Macrophage (Lyve1+), with Itgax+ and Mrc1+ macrophages in the LP and Lyve1+ macrophages in the submucosa. In disease, we observed a substantial increase in Monocytes and the emergence of Macrophage (Cxcl10+), mainly found in the LP. We also identified several subsets of dendritic cells (DC), present in both health and disease, distributed throughout the mucosa, and including conventional DC of type 1 (cDC1), DC (Fscn1+), and DC (Ccl22+). We distinguished two populations of Neutrophils. Neutrophil 1 was found in the LP, and Neutrophil 2 primarily in the lumen of inflamed tissue. We also observed multiple T cell populations, including cytotoxic [T (Cd8+)], helper [T (Cd4+, Ccr7+)], regulatory [Treg], cycling [T (Mki67+)] and follicular-helper cells [Tfh]. T cells were generally found within the LP, with Tfh enriched, as expected, in follicles. We observed natural killer (NK) cells, innate lymphoid cells of type- 2 (ILC2) and of type-3 (ILC3) with the latter mixed with lymphoid-tissue-inducer-(LTi)-like cells, all distributed largely in the LP. We identified multiple populations of B cells: B cell 1 and B cell 2 (found in the LP and within isolated lymphoid follicles), B (Aicda+) found exclusively in the dark zone of follicles, and plasma cells (found in the LP). Finally, we identified a rare population of Mast cells.

We also identified a diversity of fibroblast populations with distinct spatial locations (**Figures 1M-1P**). As there is not yet a consensus regarding the subclassification or naming of gut fibroblast populations, we adopted a nomenclature in which fibroblasts are numbered, separately for those present in healthy tissues or induced during inflammation. Nonetheless, by leveraging gene expression and spatial location, we found clear associations between previously described fibroblast groups^9–12, 15, 17, 18, 39, 40, 64, 65^ and those found here (**Figure 1M-1N; Figure S1E-S1K**). In the healthy mucosa, Fibro 6 was enriched at the base of the crypt, and its expression of *Wnt2b, Col15a1, Cd34* and *Grem1* linked this cluster to fibroblasts previously described as crypt-base fibroblasts or *Pdgfra*- low fibroblasts. Fibro 2 was enriched towards the crypt tip with clear gradients in gene expression along the crypt axis (**Figure S3Q-S3T**), and its expression of *Adamdec1, Wnt5a, Bmp2, Bmp5, Bmp7* and *Pdgfra* assigned this cluster to fibroblasts previously termed top-crypt fibroblasts or telocytes. Finally, Fibro 13 was distributed throughout the LP, and its expression of *Tagln*, *Gli1*, and a variety of integrins marked this population as myofibroblasts. Outside the mucosa, Fibro 7 was enriched in the submucosa, and its expression of *Pi16, Ackr4, Grem1, Cd34* and *C3* linked this cluster with populations previously called trophocytes, Pi16+ fibroblasts, or Ackr4+ fibroblasts. Fibro 15 was found between the muscle layers, and its expression of *Jam2*, *Ngf*, *Gdf10*, and high levels of *Pdgfra* associated it with a recently described non-mucosal fibroblast of unknown location^18^. Given its proximity to the myenteric plexus and its expression of a variety of nerve growth factors (*Ngf* and *Gdf10*), we propose Fibro 15 may play a role in ENS homeostasis. Fibro 1 was found in a single-cell layer near the serosal boundary, Fibro 4—rare in healthy tissues—was found near the edges of follicles, and Fibro 5 and 12 were found in multiple locations, although Fibro 5 was enriched in mesenteric tissue. These fibroblasts do not have clear analogs in clusters identified in previous scRNAseq studies perhaps because they are enriched in tissue regions often removed for scRNAseq (Fibro 1 and 5), are rare in healthy tissues (Fibro 4), and are marked only by a small number of genes (Fibro 5 and 12) in our library. Finally, we identified a population of fibroblast-reticular cells (FRC), which, as expected, were enriched in follicles, as well as two populations of pericytes, with Pericyte 1 enriched in the mucosa and Pericyte 2 predominantly enriched in the submucosa (**Figures 1M-1P**).

Like other cell populations, fibroblasts can also be remodeled during inflammation, giving rise to IAFs^9, 18, 39, 40^. Our measurements revealed several IAF populations with characteristic spatiotemporal distributions and gene expression profiles (**Figures 1M-1P**). IAF1 was observed at Day 3 and was marked by IEGs (e.g., *Egr1*, *Fos*, and *Nr4a1*). Based on its distribution throughout the LP and its expression of a mixture of LP fibroblast markers, IAF1 might represent a composite of the early response state of Fibro 2 and 6. IAF2 was seen at both Days 3 and 9, was marked by *Igfbp5*, *Grem1*, *Col18a1*, and low levels of *Il11*, and was found enriched at the crypt base (**Figures 1M-1P**), suggesting that it might represent a polarized form of Fibro 6. By contrast, IAF3 and IAF4 were predominantly observed only at Day 9. IAF3 was similar in gene expression and location to IAF2 but was marked by increased *Col18a1* and *Mmp2* expression, suggesting that it represents a later stage of IAF2 polarization. IAF4 emerged only at Day 9, was marked by high levels of *Il11* and *Il1β* as well as *Il1rl1*, *Mmp3*, *Mmp10*, *Mmp13*, and *Plau*, and its enrichment towards the top of crypts suggests it might represent a polarized form of a subset of Fibro 2 (**Figures 1M**–**1P**).

We also observed inflammation-induced changes in the submucosa and muscularis externa (**Figures 1M-1P**). In these regions, there were no new fibroblast populations in disease; nonetheless, there were substantial disease-associated changes. At Day 0 we observed a few rare Fibro 4 cells located mainly near lymphoid follicles. However, their number increased substantially at Day 9 and Day 21, with Fibro 4 cells found throughout the submucosal region. Given the shared markers between Fibro 7 and Fibro 4 (e.g., *Pi16*, *C3*, *Cd34*, and *Cd248*), the shared submucosal location, and the corresponding decrease in Fibro 7 abundance with the increase in Fibro 4 abundance in this location, we propose that Fibro 4 is a polarized form of Fibro 7. Based on the upregulation of cytokines (*Il1b*, *Il11*, *Csf3*, *Cxcl5*) and ECM-remodeling genes (e.g., *Mmp3*), Fibro 4 is in many respects an IAF, despite its very small presence in specific locations in healthy tissue. Similarly, we also observed an inflammatory response in Fibro 1, which extended from the serosa into the muscularis externa and submucosal regions at Day 9.

Our MERFISH measurements also revealed the expected healthy populations of EC, SMC, cells of the ENS, ICCs, and adipose cells (**Figure S3A-S3P**) and, interestingly, the emergence of novel populations during DSS colitis. For example, notable changes at Day 9 included the emergence of multiple inflammation associated SMCs (IASMCs); and the polarization of multiple neurons and glial populations, including Nos1+ neurons, Gfap+ glia, and Apod+ glia. At Day 9, we also noted the expansion of a population of lymphatic ECs expressing large amounts of the chemokine *Ccl21a*. Taken together, our MERFISH measurements reveal the extent of tissue remodeling in response to DSS-induced inflammation, highlighting contributions from nearly all cell classes in the tissue.

### Cellular neighborhoods capture DSS-induced spatial remodeling

Having charted the cellular remodeling that occurs throughout DSS-induced colitis, we next sought to define the spatial organization of the gut and its remodeling in disease. One way to quantify spatial organization is to measure cell abundance in pre-defined anatomical locations. This approach is commonplace in the context of the brain^51, 52^, where anatomical coordinates have been well established. However, we noted that while repetitive anatomical structures exist in the healthy gut, some such structures are lost in the context of disease, challenging the use of an anatomical-coordinate-based analysis (**Figure 2A**).

**Figure 2:**
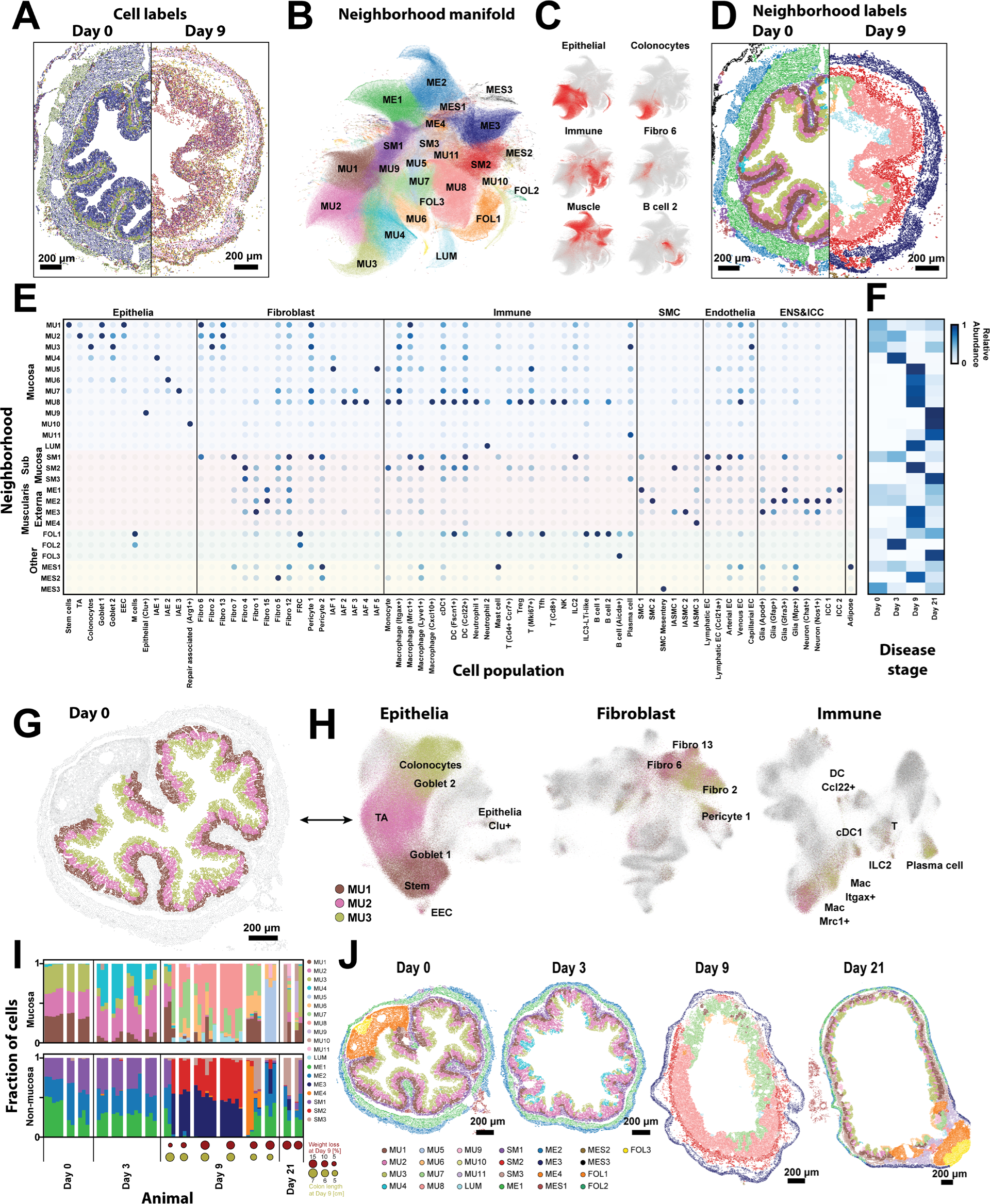
Cellular neighborhoods define the spatial remodeling of the colon during DSS-treatment. (A) The spatial distribution of all cell types in representative slices from Day 0 and Day 9. Dots represent cell centroid, and color indicates cell identity. Scale bars: 200 µm. (B) UMAP representation of the cellular neighborhoods that arise in all slices across all disease stages. Markers represent individual cells colored by the statistically distinct neighborhoods to which they were assigned. MU: mucosal; LUM: luminal; SM: submucosal; ME: muscularis externa; FOL: follicle; MES: mesentery. (C) UMAP representation of cellular neighborhoods in which each cell is colored by the fraction of its neighbors that are the listed cell class (left) or cell cluster (right). (D) The spatial distributions of cell types in the same slices in (A) colored by the tissue neighborhood to which they were assigned as in (B). (E) The fraction of each cell population found within each cellular neighborhood, normalized to the maximum value. (F) The average fraction of cells per slice assigned to each neighborhood for each disease stage, normalized by the maximum fraction observed at any day. (G) The spatial distribution of cells within a representative slice taken from Day 0 with all cells colored gray and cells assigned to each of the healthy mucosal neighborhoods, MU1, MU2, and MU3, colored as designated. (H) UMAP representation of all epithelial, immune, and fibroblast cells. Cells are colored by the neighborhood to which they have been assigned using the same colors as in (G) with all other cells not in MU1, MU2, and MU3 colored gray. (I) The fraction of cells in mucosal (top) or non-mucosal (bottom) neighborhoods measured within individual slices, grouped by the mouse of origin and the disease stage. Circles at Day 9 and Day 21 represent Day 9 measurements of the weight loss relative to Day 0 (red) or the colon length (yellow). (J) The spatial organization of all neighborhoods in representative slices taken from mice at each disease stage. Markers represent cell centroids and are colored based on neighborhood assignment. Scale bars: 200 µm.

To address this challenge, we developed an anatomical-coordinate-free statistical method for representing spatial structure. In an approach similar to that proposed by others^66, 67^, we identified statistically recurrent collections of cells, which we term cellular neighborhoods, and used these emergent neighborhoods to quantify the reproducibility of spatial organization across biological replicates, the development of different neighborhoods across disease, and the organization of such neighborhoods with respect to anatomical features within the tissue. We identified 25 cellular neighborhoods (**Figures 2A-2D**), collectively comprising 99.5% of all cells, each with characteristic cell compositions (**Figure 2E**), locations (**Figure S4**), and prominence in different disease stages (**Figure 2F**). We named neighborhoods based on their anatomical location and numbered them in the order of emergence throughout DSS-disease course, producing neighborhoods associated with the mucosa (MU1-11), lumen (LUM), submucosa (SM1- 3), muscularis externa (ME1-4), follicles (FOL1-3), and mesentery (MES1-3).

We initially investigated the healthy mucosal neighborhoods to validate our analysis (**Figures 2E-2H**). In homeostasis, the maturation of epithelial cells from intestinal stem cells to mature colonocytes is orchestrated, in part, by a fibroblast-derived Wnt-Bmp gradient along the crypt^68^. Indeed, our healthy mucosal neighborhoods, MU1-3, recapitulate this known zonation (**Figures 2G-2H**). MU1, found at the crypt base, was enriched in intestinal stem cells and Fibro 6 which produced notable stem-cell maintenance factors (e.g., *Wnt2b* and *Grem1*); MU3, found at the crypt tip, was enriched in colonocytes and Fibro 2 which expressed top-crypt epithelial maturation factors (e.g., *Bmp2*, *Bmp5*, and *Bmp7*); and, MU2, found in the middle of the crypt, was enriched in TA and a mixture of Fibro 2 and 6, reflecting this intermediate stage in the Wnt-Bmp gradient (**Figures 2G-2H**; **Figure 1N**). In addition, macrophages shift between more phagocytic to bactericidal (Mrc1+ or Itgax+) phenotypes as they migrate from the crypt base to the tip in a microbiota-dependent process^69, 70^. Our neighborhoods reproduced this distribution as well, with MU1 enriched in Mrc1+ macrophages, MU3 enriched in Itgax+ macrophages, and MU2 containing intermediate proportions of both (**Figure 2H**).

Having provided support for this analysis, we next leveraged these neighborhoods to explore the uniformity of tissue response to DSS treatment. Neighborhood composition was highly reproducible in healthy mice in both the mucosal and non-mucosal regions, underscoring the regular structure of the healthy gut (**Figure 2I**). By contrast, at Day 3, the inter-slice variation increased in the mucosal neighborhoods but remained similar to that observed at Day 0 for the non-mucosal neighborhoods, reflecting an earlier inflammatory response of the mucosa relative to the non-mucosal gut regions (**Figure 2I**). At Day 9, both regions of the gut had substantial inter-slice and inter-mouse variation, revealing a remarkably heterogeneous tissue response (**Figure 2I**). Interestingly, even after the mice had regained their pre-treatment weight at Day 21, there was still substantial variation between slices (**Figure 2I**), indicating that the variability induced during disease persists during repair.

This variability was also apparent in space (**Figure 2J**; **Figure S4**). At Day 0, the organization of the mucosa into three mucosal neighborhoods was highly conserved from crypt to crypt both within and between mice. By contrast, at Days 3, 9, and 21, this highly regular organization was largely disrupted, and contiguous regions of the mucosa, often spanning the length-scale associated with multiple adjacent crypts, were assigned to different neighborhoods (**Figure 2J**), revealing both a spatially variable and a spatially coordinated response at the scale of adjacent crypts. A non-uniform or ‘patchy’ response to DSS-induced disease or IBD has been reported previously though typically discussed in the context of histology^44, 45^. Complementing this previous work, our analysis now reveals the molecular and cellular correlates of this spatial variability.

The phenotypic response of genetically identical co-housed mice subject to DSS-colitis is known to be variable^71^. Thus, we asked to what degree the variation in disease severity— as measured by weight loss and colon shortening—might correlate with the variability in neighborhood usage (**Figure 2I**). Suggestively, the mouse with the least severe disease measures at Day 9 had an increased frequency of healthy neighborhoods—MU1, MU2, and MU3—while the mice with more severe measures tended to have greater proportions of MU7, MU8, and LUM, raising the possibility that disease severity is, indeed, linked to neighborhood usage. Nonetheless, given the variability in neighborhood usage and phenotype, a link between these correlates would require further study.

### Gut remodeling occurs via a staged progression of cellular neighborhoods

To better understand multiple features of these neighborhoods, including their cellular composition, spatial relationship to one another, and potential transformation from one neighborhood to the next throughout disease, we investigated the neighborhoods in greater depth. Starting with the mucosa, we noted that at Day 3 the tissue organization was largely unchanged from Day 0. Nonetheless, our neighborhood analysis revealed a variety of subtle cellular and molecular changes (**Figure 3A**). First, there was a reduction in the abundance of MU1, reflecting an overall decrease in stem cell numbers per slice (**Figure 1G**). Interestingly, this change was not mirrored by a similar reduction in the number of Fibro 6 (**Figure 1O**) or their expression of the key stem-cell trophic factors, e.g., *Wnt2b* (**Figure S5A**), suggesting a stem-cell intrinsic DSS response. Where MU1 was lost, stem cells were replaced with TA cells, further reflecting the aging of the epithelia, and MU1 was generally replaced by MU2 (**Figure S5B**). Moreover, even where they were preserved, MU1, MU2, and MU3 had subtle shifts in cell-type composition relative to Day 0: most notably, an increase in macrophages with more Mrc1+ macrophages in MU1 and MU2 and more Itgax+ macrophages in MU3 (**Figure S5C**).

**Figure 3:**
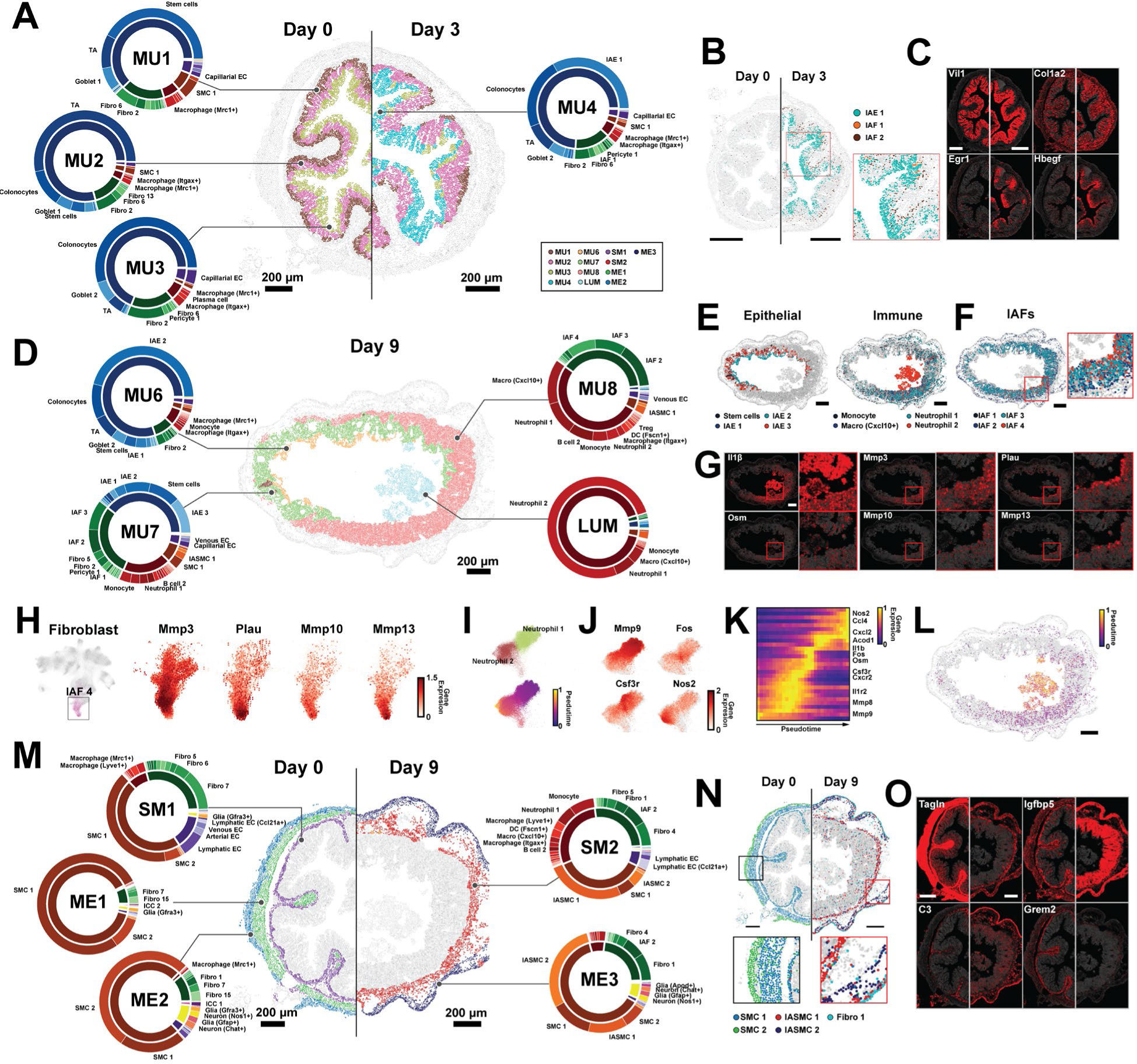
Cellular neighborhoods evolve in a staged progression driven by polarization and recruitment of distinct cell populations. (A) The spatial distribution of mucosal neighborhoods in a representative healthy (Day 0) and early inflammation (Day 3) slice. Markers represent the cell centroids and color indicates neighborhood assignment. Pie charts represent neighborhood composition of cell classes (inner; fibroblast: green, epithelial: blue, immune: red, EC: purple, SMC: brown, ENS: yellow) or cell clusters (outer). Prominent populations are labeled. Scale bars: 200 µm. (B) The spatial distribution of cell populations that define MU4 in representative Day 0 or Day 3 slices. Inset: zoom-in on red boxed region. Scale bars: 200 µm. (C) The spatial distribution of RNAs marking populations that define MU4 in the slices in (B). The centroids of all RNAs are plotted in gray and the listed RNAs in red. Scale bars: 200 µm. (D) As in (A) but for mucosal neighborhoods that emerge at Day 9. Scale bar: 200 µm. (E) The spatial distribution of cell populations that define Day 9 neighborhoods plotted as in (B). Scale bars: 200 µm. (F) The spatial distribution of IAFs highlighting variation with the MU8 ulcer. (G) RNA spatial distributions for key markers of Day 9 neighborhood populations as in (C) for the slice in (D) with zoom in on the red boxed regions. Scale bar: 200 µm. (H) UMAP representation of fibroblasts (left) with IAF4 cells colored and a zoom in on the IAF4 region of this UMAP with individual cells colored by genes that define IAF4 and IAF4 subdivisions (right). (I) Zoom-in on the portion of the immune UMAP that shows Neutrophil 1 and Neutrophil 2 with cells colored by cell population (top) or pseudo time (bottom). (J) Zoom-in on immune UMAP highlighting gene expression within Neutrophil 1 and Neutrophil 2. (K) The normalized expression of highlighted genes sorted by neutrophil pseudo time as in (I). (L) The spatial distribution of Neutrophil 1 and Neutrophil 2 colored by pseudo time for the slice in (D). Scale bar: 200 µm. (M) As in (A) but for non-mucosal neighborhoods in Day 0 and Day 9. Scale bars: 200 µm. (N) As in (B) but for cell populations that define Day 0 and Day 9 non-mucosal neighborhoods with black and red boxed zoom-in (bottom). Scale bars: 200 µm. (O) As in (C) but for RNAs that mark cell populations in highlighted in (N).

In addition, a defining feature of the Day 3 mucosal response was the emergence of MU4. MU4 was characterized by three inflammation-associated populations—IAE1, IAF1, and a small amount of IAF2—and its extent was shaped by the distribution of these cells and the genes they express (**Figures 3A-3C**; **Figure S5B**). The observation that IAE1 and IAF1 predominantly co-occur in the same neighborhood suggests that the polarization of these two populations is spatially coordinated. In contrast to the remodeling that occurred in the mucosa at Day 3, no new neighborhoods arose in non-mucosal regions, suggesting that the mucosa is the first region to respond to DSS.

At Day 9, the mucosa was almost completely remodeled and was largely defined by four emergent neighborhoods, MU6, MU7, MU8, and LUM, with some re-emergence of MU1 (**Figures 3D-3G**). The recovery of MU1 was driven by an overall increase in the number of stem cells (**Figure 1G**), suggesting that stem cell recovery begins rapidly after DSS withdrawal (**Figure 1A**). Nonetheless, MU6, MU7, and MU8 represented 75% of mucosal cells at Day 9. The crypt base-to-tip zonation was largely abolished on Day 9, and MU6 and MU7 tended to co-occur, with MU6 almost exclusively at the tip of flatted crypts and MU7 or MU1 below it (**Figure 3D**; **Figure S4**). MU8, by contrast, was observed either under regions of MU7 in which MU6 had been lost or, more frequently, across the entire mucosal region (**Figure 3D**; **Figure S4**).

MU6 was similar in composition to MU3 but distinguished, in part, by the emergence of IAE2 (**Figures 3D**-**3E**), which given its pro-inflammatory, anti-bacterial signature (**Figure 1F**) suggests that MU6 is a pro-inflammatory version of the top-crypt, MU3 neighborhood. MU7 was characterized by distorted, hyperplasic crypts (**Figure 3D**; **Figure S4**), distinguished based on the abundance of stem cells and stem-like IAE3, suggesting that MU7 corresponds to a crypt neighborhood with rapid, dysregulated epithelial proliferation. This observation, in turn, suggests a molecular signature for hyperplasic crypts (**Figures 1F** and **3D-3F**). In addition, MU7 was defined by a reduction in healthy goblet cells, the presence of multiple IAF populations—IAF1, IAF2, and IAF3—and a dramatic increase in the abundance of multiple of immune populations, including those less abundant in early stages of diseases, such as Monocytes and Neutrophil 1 (**Figure 3D-3F**).

Another prominent feature of Day 9 tissue was MU8 (**Figure 3D**). MU8 was distinguished by an almost complete lack of an epithelial layer, marking this region as ulcerated tissue (**Figure 3D-3E**). MU8 was further defined by the presence of IAF4, in addition to IAF2 and IAF3, and an even greater enrichment and diversity of immune populations as that seen in MU7. Indeed, roughly 50% of cells within this neighborhood were immune cells (**Figure 3D-3E**). Of these, Macrophage (Cxcl10+), Neutrophil 1, and B cell 2 were the most prominent, though we also observed notable amounts of Monocytes, Neutrophil 2, DC populations (cDC1, DC [Ccl22+] and DC [Fscn1+]), ILC3-LTi-like cells, ILC2, NK cells, Tregs, T (Cd4+, Ccr7+), and T (Cd8+) (**Figures 3D** and S5D**-S5E**). The absence of healthy fibroblasts combined with the enrichment in IAFs and immune populations underscores the highly inflammatory environment of this neighborhood.

In addition, we noted several sub-structures within the MU8 ulcers. First, IAF4 was enriched at the luminal surface while IAF2 and IAF3 were depleted in this region. Furthermore, we noted gene expression gradients within IAF4 itself, with several genes, such as *Mmp13*, *Mmp10*, and *Plau*, expressed at high levels in IAF4 cells closer to the luminal surface (**Figure 3G-3H**). Given the differential expression of matrix metalloproteinases (Mmp) between IAFs and within IAF4, this observation suggests that a spatial fine tuning of ECM properties within the MU8 ulcers is an important function of IAFs. Second, we observed an enrichment in Neutrophil 2 and Macrophage (Cxcl10+) near the luminal surface, DC (Fscn1+) near the muscularis mucosa, and an enhanced B cell density in small sub-regions of MU8 near the muscularis mucosa (**Figures S5D**-**S5E**), raising the possibility of compartmentalized immune responses within the ulcerated tissue.

Finally, MU8 showed evidence of vascular remodeling, with the emergence of venous ECs, which were not seen in the Day 0 or Day 3 mucosa (**Figure 3A**). Interestingly these venous EC were organized into discrete semi-regular clusters (**Figure S5F**), suggesting that while the overall crypt structure is lost in MU8, some tissue regularity remains. Relative to venous EC in healthy tissues, the MU8 venous ECs upregulated a variety of immune-recruitment genes—e.g., *Selp* and *Cxcl10*—and downregulated the EC tight junction gene, *Cldn5*, underscoring both the increased vascularization of the MU8 ulcers and a functional tuning of these cells likely important for immune recruitment to MU8 (**Figure S5F-S5H**).

Adjacent to MU8 in many slices was the LUM neighborhood (**Figure 3D**; **Figure S4**). This neighborhood was composed mainly of Neutrophil 2 (**Figure 3D**). As Neutrophil 2 was also found in MU8, we asked if Neutrophil 2 might be a polarized form of Neutrophil 1 excreted into the lumen from MU8. Supporting this notion, we noted that a subset of both Neutrophil 1 and Neutrophil 2 expressed a set of IEGs, consistent with a transcriptional intermediate associated with this possible polarization (**Figure 3I-3J**). To formalize this observation and identify additional transition genes, we used a pseudo time analysis to sort cells along the possible transition from Neutrophil 1 to Neutrophil 2 (**Figure 3I-3K**). In addition to multiple IEGs, this analysis identified several genes enriched at intermediate pseudo times (**Figure 3K**), most notably *Osm*—a recently identified biomarker of IBD- severity for which neutrophils in ulcerated tissue are a major source^16, 72^. Our measurements now suggest that *Osm* may be produced by neutrophils primarily as they transition from mucosal (Neutrophil 1) to intra-luminal (Neutrophil 2). We also noted that transitioning neutrophils did not have a clear spatial organization within MU8, suggesting that this transition is not tied to interactions in any specific subregion (**Figure 3L**).

Unlike in Day 3, in Day 9 we observed changes to non-mucosal neighborhoods, with a thickening of the muscularis mucosa, muscularis externa, and submucosa (**Figure 3M; Figure S4**). Our measurements revealed that these expected anatomical changes^44^ are reflected in changes in cell composition and the emergence of new neighborhoods for these regions. At Days 0 and 3, SM1 represented the submucosa and muscularis mucosa and was defined by Fibro 7, ECs associated with blood or lymphatic vasculature, and SMC1 associated with the muscularis mucosa. At Day 9, this region was defined by a new neighborhood, SM2, which, in turn was defined by the replacement of Fibro 7 with Fibro 4, lymphatic EC with Ccl21+ lymphatic EC, SMC1 with IASMC1, and an increased abundance of Monocytes, Neutrophil 1, DC (Fscn1+), and B cell 2 (**Figure 3M**). We also noted subtle changes in gene expression within some cell populations in SM2. Specifically, relative to those in SM1, Lyve1+ macrophages in SM2 had statistically significant enrichment in chemokines that attract monocytes (Ccl8) and B/T cells (Cxcr5) (**Figure S5I**). As Ccl21 is also a chemoattractant for a variety of immune populations, these shifts support a coordinated effort for immune recruitment in SM2 and the mucosal layer above. Inflammatory responses in the muscle layers are less well described in the context of colitis models; yet at Day 9, the emergence of multiple IASMC populations led to the replacement of ME1 and ME2 with a single neighborhood, ME3, that captured both layers. Interestingly, ME3 was also defined by an increased abundance of Fibro 1, which reflected the inflammation-associated polarization of this population.

While our measurements represent static snapshots at distinct stages in the evolution of DSS response, we noted that the shared anatomical locations and overlap in cell populations that comprise different cellular neighborhoods suggested a temporal evolution to these neighborhoods. The non-mucosal response was straightforward: ME1 and ME2, seen at Day 0 and Day 3, collectively polarize to become ME3 at Day 9 while the submucosal region, SM1, polarizes to SM2 at Day 9. A similar connection was suggested with the mucosal neighborhoods though complicated by mucosal zonation. We propose the following staged polarization of the mucosa. Healthy MU1, MU2, and MU3 polarize to MU4 in early inflammation; MU4 transforms to MU6 and MU7 during peak inflammation; MU6 is lost while MU7 transforms to MU8; and MU8 produces LUM.

Thus, taken together our neighborhood analysis revealed a broad cellular and spatial remodeling in all regions of the mouse colon in response to DSS-induced damage with the spatial response described by a series of statistically distinct cellular collections— neighborhoods—with defined cell compositions, sharp spatial boundaries, and coordinated spatial responses. These neighborhoods, in turn, likely represent a staged evolution of the tissue from the organization seen in healthy tissues to the late-stage inflammatory regions seen at Day 9.

### Recovery associated neighborhoods emerge after DSS removal

At Day 21, two weeks after DSS withdrawal, most mice had recovered their pretreatment body weight. Thus, we leveraged our neighborhood analysis to assess to what degree this phenotypic recovery was reflected in the molecular and cellular landscape of the gut. In the mucosa at Day 21, we observed regions of the tissue in which the healthy neighborhoods, MU1, MU2, and MU3 re-emerged (**Figure 4A**). However, these neighborhoods did not reflect the healthy crypt structure seen at Day 0. Moreover, they had subtle shifts in cell-type abundance relative to Day 0, most notably a marked increase in the abundance of plasma cells (**Figure S5C**) which likely reflects the established, long-lived adaptive immune response to DSS treatment^39, 73^ and may mirror a similar phenomenon, known as basal plasmacytosis, in human IBD.

**Figure 4:**
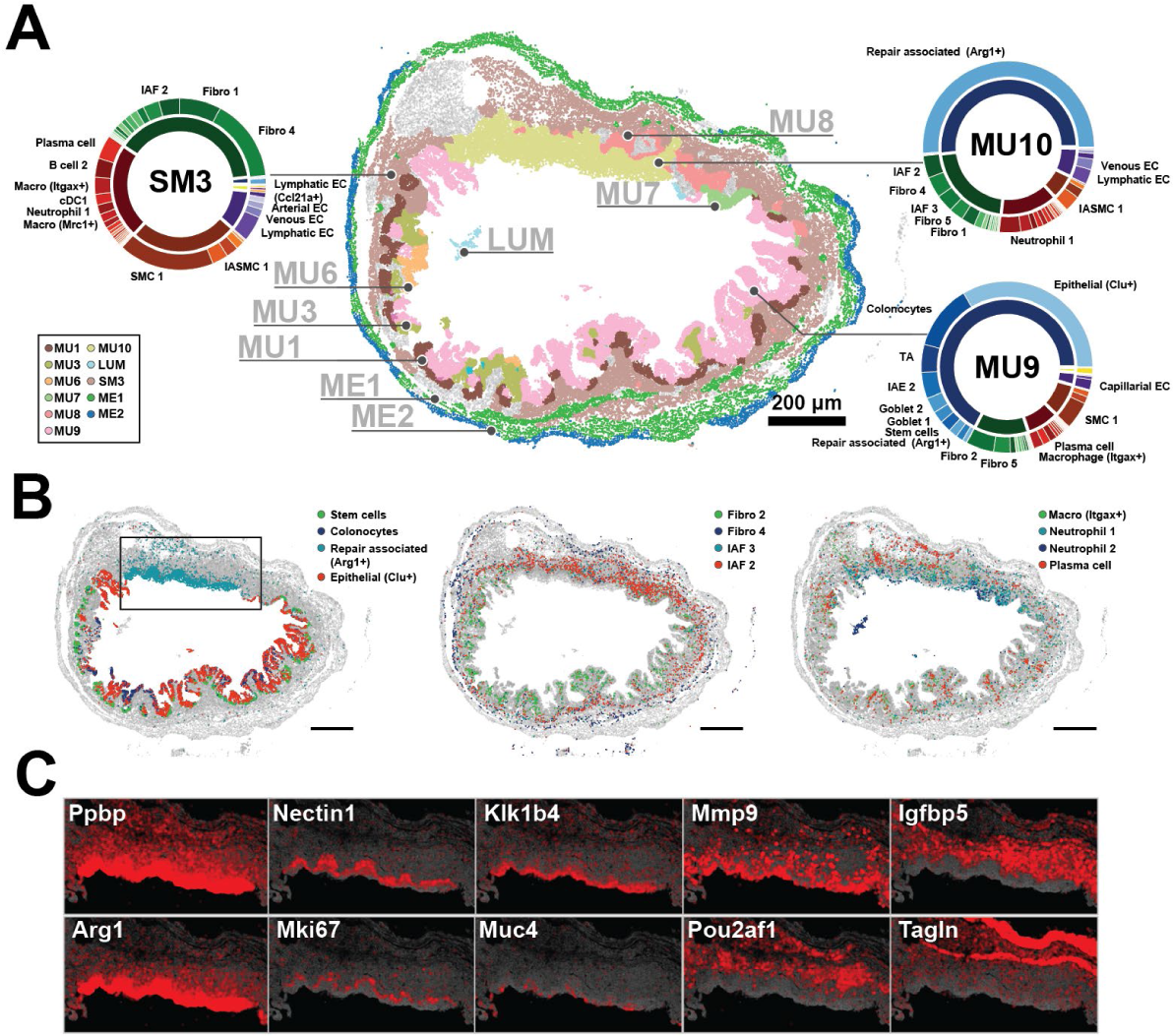
Cellular neighborhoods define the organization of repairing tissue. (A) The spatial distribution of mucosal neighborhoods in a representative repairing slice (Day 21). Markers represent the cell centroids and color indicates neighborhood assignment. Pie charts represent neighborhood composition of cell classes (inner) or cell clusters (outer). Prominent populations are labeled. Scale bar: 200 µm. (B) The spatial distribution of cell populations that define neighborhoods observed at Day 21. Scale bars: 200 µm. (C) The spatial distribution of RNAs that highlight the structure of the MU10 repairing ulcer plotted for the black boxed region in (B). The centroids of all RNAs are plotted in gray and the listed RNAs in red.

In addition, we observed the emergence of several novel mucosal neighborhoods seen only in Day 21—MU9, MU10, and MU11—and these neighborhoods were often intermixed with regions of MU1, MU2, and MU3 as well as some inflammatory neighborhoods such as MU6, MU7, and MU8 (**Figure 4A; Figure S4**). MU9 was associated with a partial restoration of the crypt structure and healthy epithelial populations, such as stem, TA, colonocytes, and both goblet cell clusters (**Figure 4A-4B**). However, MU9 was distinguished by a substantial number of Clu+ epithelial cells, the absence of IAFs, and the presence of Fibro 2 but little Fibro 6 or Fibro 13 (**Figure 4A- 4B**), suggesting that MU9 represents an intermediate stage in the repair of the healthy mucosa.

MU10 had no evidence of crypt structure, contained some IAFs, was enriched in immune populations, and was the dominant source of Repair-associated epithelia, marking this neighborhood as a repairing ulcer (**Figure 4**). Just as we observed substructure within the MU8 ulcer, we also noted an intricate spatial organization within the MU10 repairing ulcer. Specifically, we observed strong gene expression gradients within the repair-associated cells, suggesting multiple functionally distinct layers of these squamous epithelia (**Figure 4C**). Similarly, distinct immune populations were enriched in different regions of MU10, with Neutrophil 2 enriched near the luminal surface, Neutrophil 1 distributed throughout, and a layer of plasma cells found near the base of MU10 (**Figure 4B**). Finally, we observed a few rare instances of MU11, which was found near the base of the mucosa and was defined almost entirely of B cell 2 and plasma cells (**Figure S4; Table S3**).

Outside the mucosa we largely observed restoration of the healthy muscularis externa neighborhoods, ME1 and ME2, indicating partial recovery from the inflammation associated changes seen in Day 9 (**Figure 4A**; **Figure S4**). By contrast, the muscularis mucosa and the surrounding submucosa were defined by a novel neighborhood, SM3, which shared the SMC usage with that of healthy SM1 and the immune and fibroblast usage with that of SM2 (**Figures 3M** and **4A**). The only notable difference between the immune populations in SM3 and SM2 was an increase in the abundance of Plasma and B cell 2 in SM3 (**Figures 3M** and **4A**). Overall, the partial recovery of the submucosa is consistent with the observation that SMCs recover from inflammation-induced states more rapidly than fibroblast and immune populations. Taken together, these observations demonstrate that even when mice have regained their pre-treatment body weight, the cellular and molecular structure is far from that observed in the healthy gut. Instead, the tissue is comprised by a series of emergent neighborhoods that either represent different stages of repair or the differential repair of various Day 9 neighborhoods.

### Inflammation-associated fibroblasts orchestrate unique cell-cell interactions

We next asked what modes of interaction might play critical roles in shaping the composition, development, and cellular behaviors within these neighborhoods. Receptor-ligand analysis tools have proven powerful for proposing rich interaction hypothesis sets from single-cell measurements^74–76^. However, since the strength of many interactions are modulated by physical proximity, the lack of spatial context in these measurements challenges the prioritization of proposed interactions. To address this challenge, we developed a receptor-ligand analysis that leverages cellular co-occurrence within cellular neighborhoods to provide a set of spatially prioritized hypotheses for cell-cell interactions. We adopted this approach as opposed to recent similar methods that leverage direct cell proximity^77^, as we reasoned that neighborhood co-occurrence might better capture meaningful interactions not as easily revealed by direct cell-cell contact, such as signaling via diffusible ligands or transient interactions between mobile cell populations. Application of this approach produced a rich set of interaction hypotheses for each of our identified neighborhoods.

To provide some support for these proposed interactions, we first focused on the healthy mucosa (**Figures 5A-5B**) and identified a spectrum of well-established communication modes. These validated interactions included the trophic interactions^68^ between epithelial stem cells expressing Wnt receptors (*Lrp5/6* and *Fzd1-6*) with Wnt ligands (*Wnt2b* and *Wnt4*) expressed by Fibro 6 (**Figure 5A**); the recruitment of macrophages^78^ expressing the chemokine receptor *Cx3cr1* to the LP by epithelial cells expressing *Cx3cl1* (**Figure 5B**), and the interaction between macrophages^79^ expressing *Csf1r* and the various healthy LP fibroblasts, Fibro 2, 6, and 13, all of which express *Csf1* (**Figure 5B**).

**Figure 5:**
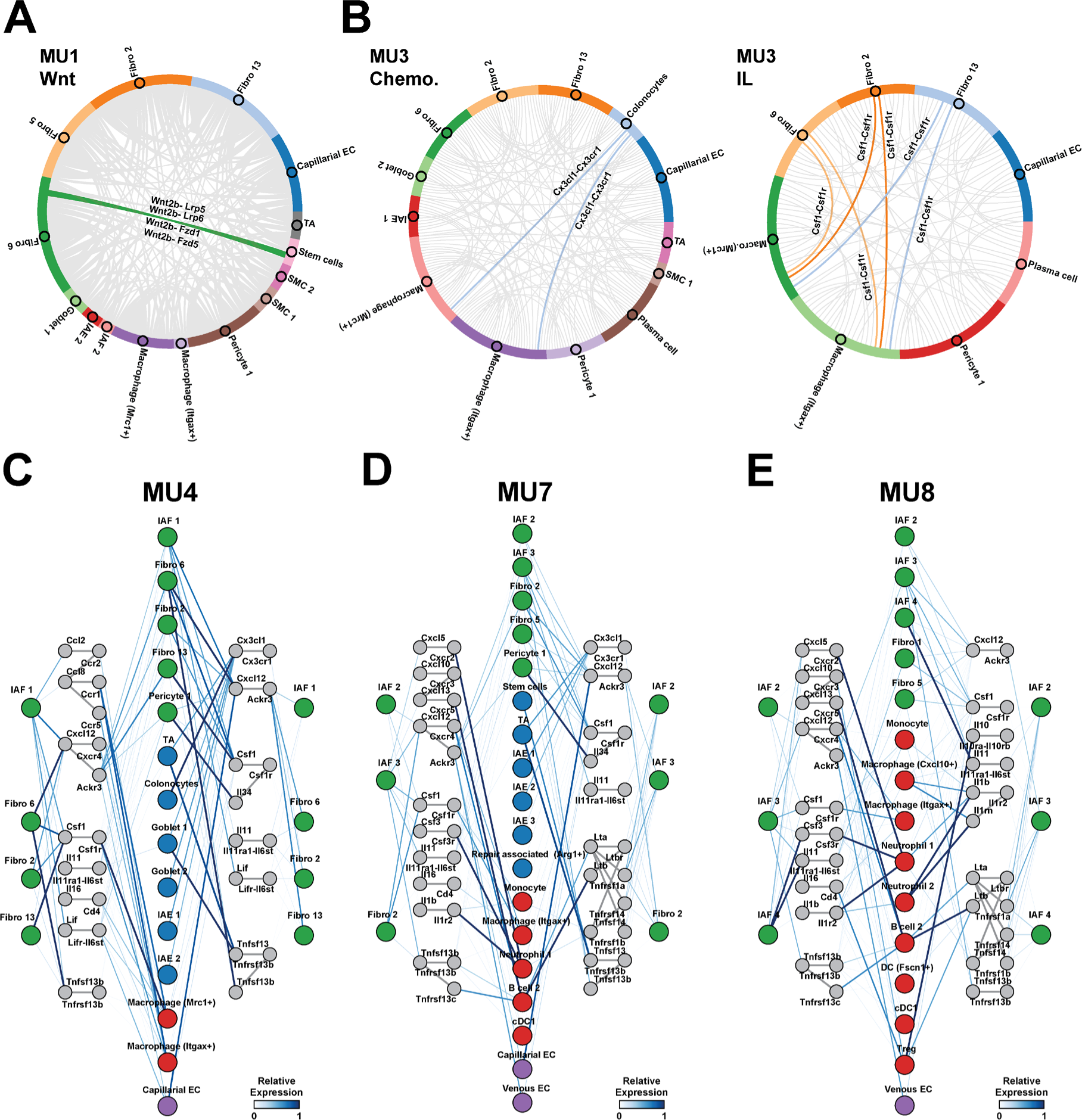
Spatially prioritized receptor-ligand interactions suggest unique roles for IAFs in different cellular neighborhoods. (A) Circos plots showing the interaction frequency between cell populations in MU1 for all Wnt-family ligand-receptor interactions. Highlighted connections represent key known fibroblast-epithelial interactions. (B) As in (A) but for chemokines (left) and interleukins (right) for all cell populations in MU3. Highlighted connections represent key known fibroblast-immune interactions. (C-E) Putative interaction map describing signals to and from fibroblasts in inflammation-associated neighborhoods, MU4 (C), MU7 (D), and MU8 (E). Colored circles represent major cell populations in these neighborhoods with color indicating cell class (fibroblast: green, epithelial: blue, immune: red, EC: purple). Gray colored circles represent ligands or receptor complexes and lines connecting these circles represent documented interacting pairs. Lines connecting cell populations to ligands or receptors represent expression in those populations and colored by the average expression within that population normalized to the maximum observed in any population in any neighborhood.

Given the emerging role of mesenchymal-immune interactions in orchestrating key tissue responses to inflammation^3, 8^ and the increased diversity of IAFs that we observed here relative to that previously reported^9, 15, 18, 39, 40^, we next chose to focus in depth on the cytokine-mediated interactions that IAFs might leverage to shape the composition of neighborhoods in which they are found and which, in turn, might contribute to their polarization from healthy fibroblasts. IAF1, IAF2, IAF3, and IAF4 were predominantly found within three mucosal neighborhoods—MU4, MU7, and MU8—and our receptor-ligand analysis predicted an intricate web of interactions between all fibroblasts in these neighborhoods—both healthy and IAF—and other cell populations (**Figure 5C-5E**).

We noted both common fibroblast interaction modes as well as those unique to IAFs and IAF subsets (**Figure 5C-5E**). For example, our analysis highlighted the common role of fibroblasts—healthy and IAF—in the recruitment and maintenance of macrophages via the *Cxc12-Cxcr4* and *Csf1-Csf1r* axes, and that there may be specialization or fine tuning of these interactions within different fibroblast subsets. One notable example of this fine tuning was IAF1, which was the only IAF to express significant levels of *Ccl2* and *Ccl8*, which may reinforce the recruitment of monocytes and macrophages expressing *Ccr1*, *Ccr2*, and *Ccr5* early in disease (**Figure 5C**). IAFs were also involved in the interaction with other immune populations (**Figure 5D-5E**). For example, IAF2, IAF3, and IAF4 likely aid in the recruitment of Tregs and cDC1 to MU8 via the *Cxcl10-Cxcr3* axis. Similarly, IAF3 might be involved in B cell 2 recruitment via the *Cxcl13-Cxcr5* axis while IAF2 might send signals to Tregs via the *Tnfs9-Tnfrsf9* axis and receive signals via the *Il10- Il10ra/Il10rb* axis. In addition, IAF3 and IAF4, which were the only IAFs to express a significant amount of *Csf3*, could contribute to *Csf3r*-dependent neutrophil and, perhaps, macrophage activation (**Figure 5D-5E**). Moreover, the expression of *Cxcl5* by IAF3 or the combined expression of *Cxcl1*, *Cxcl2*, and *Cxcl5* by IAF4, could contribute to MU8 Neutrophil 1 recruitment via *Cxcr2*. Finally, IAF4 may have a unique role in regulating the degree of inflammatory signaling in MU8, as it was the only IAF to express significant amounts of *Il1r2*, a decoy receptor for *Il1β*—an inflammatory cytokine abundant in MU8.

Our analysis also revealed a series of possible interactions between IAFs and all fibroblasts, both healthy and IAF, which might contribute to the activation or polarization of these fibroblasts. Notably, IAFs signaled to all other fibroblasts via *Il11* and *Lif*. As *Il11* can induce fibroblast activation in a range of contexts^80–82^, these IAF-fibroblast interactions might serve as a positive feedback mechanism to further polarize LP fibroblasts. The only exception to this observation was IAF4, which lacked the significant expression of the *Il11ra1-Il6st* receptor complex seen on all other LP fibroblasts. This observation suggests that while IAF4 may contribute to the polarization of LP fibroblasts via *Il11* expression, this may not be a relevant interaction for IAF4 polarization (**Figure 5E**). Thus, our analysis suggests that IAFs orchestrate an intricate web of interactions with immune cells and other fibroblasts, and that individual IAF subsets have specialized roles in this interaction network.

### Lineage-tracing reveals distinct origins of inflammation-associated fibroblasts

Given their distinct gene expression, spatial localization, and proposed cell-cell interactions, we next asked if different IAF subpopulations might arise from different healthy fibroblast populations—as suggested by multiple aspects of our MERFISH data. To explore this question, we leveraged a fibroblast-lineage tracing mouse generated by crossing CXCL12-creER mice with LSL-tdTomato mice to create a CXCL12^Lin^ mouse in which all cells expressing high levels of *Cxcl12* at the time of cre induction will be marked by tdTomato^83^. As Fibro 6 expresses the highest level of *Cxcl12* in the healthy LP, this model should predominantly mark Fibro 6 cells in this region. Indeed, we found that the majority of CXCL12^Lin^ cells overlapped with fibroblasts as revealed by immunofluorescence and co-expression of podoplanin (PDPN; **Figure 6A**)—a pan mesenchymal marker—but not co-expression of EC or pericyte markers (**Figure S6A- S6B**); were enriched in the sub-epithelial region at the base of the crypt, as expected for Fibro 6 (**Figure 6A**); and were enriched in a variety of genes that mark Fibro 6 (i.e., VCAM1, CTGF [CCN2], and SDC2; **Figure 6B-6D**). Flow cytometry analysis validated enrichment of CXCL12^Lin^-Tomato signal in a fraction of PDPN+ LP fibroblasts (**Figure S6C-S6D**). While we cannot rule out that the CXCL12^Lin^ mouse marks small populations of other cells, we conclude that most of Fibro 6 are marked by this lineage tracer.

**Figure 6.**
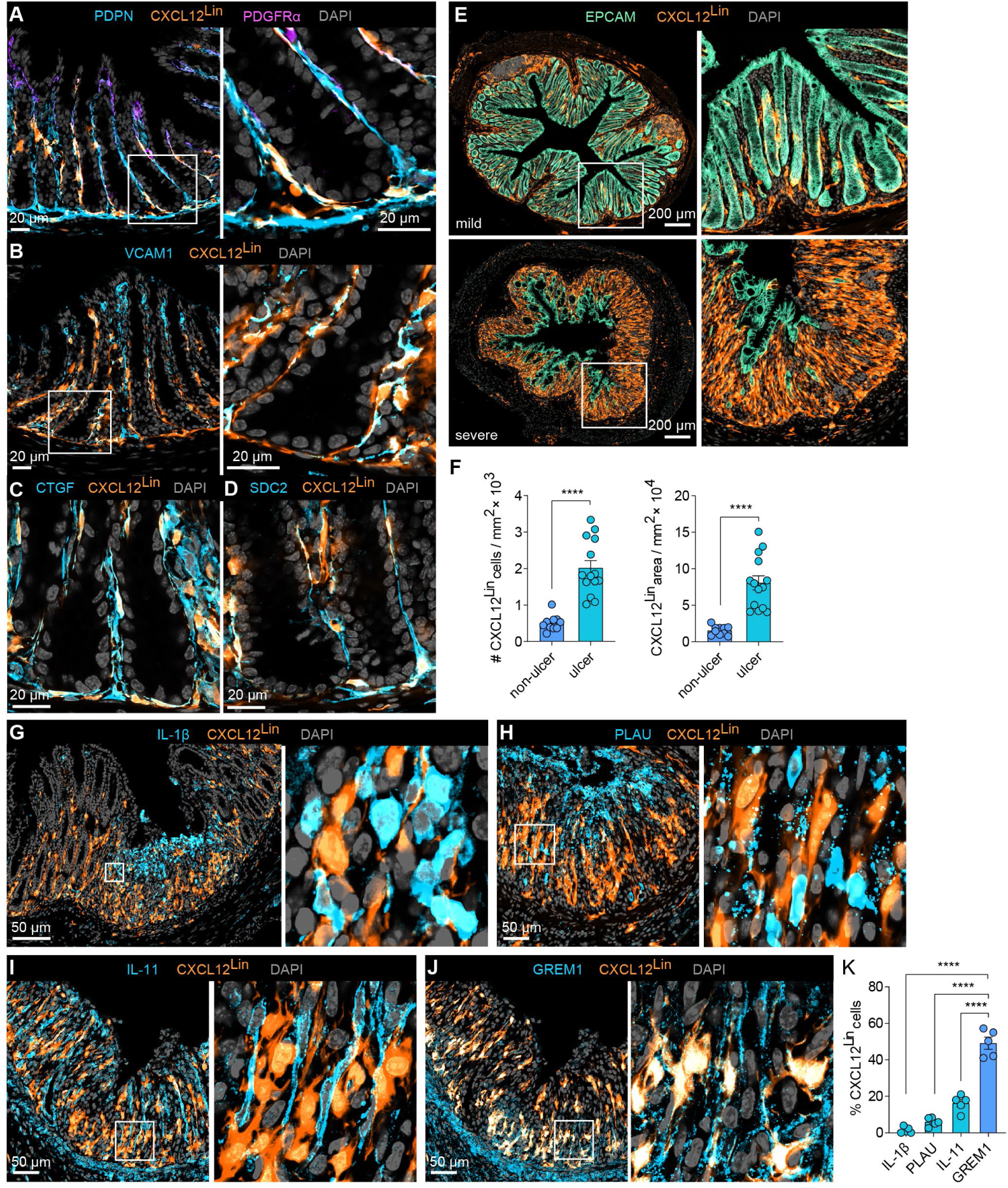
Lineage tracing reveals distinct origins of IAF populations. (A) Representative immunofluorescence images of colon cross sections from Day 0 mice (*n*=5) highlighting the overlap of CXCL12^Lin^ cells with the pan-fibroblast marker PDPN and low overlap with top-crypt PDGFRα^hi^ cells. A zoom-in on boxed region is highlighted (right). Scale bar: 20 µm. (B-D) As in (A) but for markers of Fibro 6: VCAM1 (B), CTGF (C), and SDC2 (D). Scale bars: 20 µm. (E) Representative immunofluorescence images of colon cross sections from Day 9 mice sorted by mild or severe weight loss. Right panels show magnification of boxed areas (*n*=5-6). Scale bar: 200 µm. (F) CXCL12^Lin^ cell number and area in tissue sections collected at Day 9 in ulcerated or non-ulcerated tissue regions (*n*=6-7). The bar represents the mean, the error bars represent standard error of the mean (SEM), and markers are individual mice. ****p<0.0001. (G-J) Representative immunofluorescence images of ulcerated regions, harvested on Day 9 showing overlap of CXCL12^Lin^ with IL-1β, PLAU, IL-11, or GREM1. (K) The fraction of cells positive for CXCL12^Lin^ that overlap with IL-11, GREM1, IL-1β and PLAU in ulcerated tissue regions as shown in (G-J). The bar represents mean, the error bars represent SEM, and markers are individual mice. ****p<0.0001.

To then determine the Fibro-6 origins for different IAF populations, we assessed the CXCL12^Lin^ population dynamics following DSS treatment. By segregating mice according to disease severity (**Figure S6E**), we found that CXCL12^Lin^ fibroblasts increased in abundance in regions of epithelial erosion and ulceration (**Figure 6E-6F** and **Figure S6F**), consistent with the polarization of these fibroblasts into IAFs. To then explore the overlap between different IAF populations and CXCL12^Lin^ fibroblasts, we co-stained regions of inflamed tissue with IL-1β, PLAU, IL-11 and ST2 (IL1RL1) to mark IAF4, and GREM1, VCAM1 and C3 to mark IAF2/3 (**Figures 6G-6K**; **Figure S6G-S6I**). As expected from our MERFISH measurements, IAF2/3 markers were enriched at the bottom and IAF4 markers at the top of ulcers, with some degree of intermixing. However, we observed a clear segregation in the overlap of these markers, revealing that CXCL12^Lin^ marked IAF2/3 but not IAF4 (**Figures 6G-6K; Figures S6G-S6I**). Interestingly, while CXCL12^Lin^ fibroblasts were largely excluded from the top of the ulcer, they were often in proximity with inflammatory IL-1β+ cells, including IAF4 cells (**Figures 6G-6K; Figures S6G-S6I**), suggesting potential interactions between these populations. Together our measurements reveal that the distinct IAF populations we identified with MERFISH have, in part, distinct healthy LP fibroblast origins and support a model in which the polarized form of different healthy LP fibroblasts might play distinct roles in the orchestration of tissue response to inflammation.

### Inflammation-associated fibroblast subsets and inflammation-associated neighborhoods may contribute to human ulcerative colitis

IAF populations have been identified in IBD patients, both in UC^9, 31^ and CD^30, 32, 36, 41^. We therefore next asked if the IAF populations observed here are similar to the single Il11+ IAF population seen in human UC^31^. We first noted that the human homologs of key mouse markers of the IAF populations we defined here were collectively expressed within the single Il11+ IAF population observed in humans (**Figure 7A**), suggesting that either this single IAF population serves the same functional roles as the multiple IAFs we observe in mouse or the increased throughput and sensitivity provided by MERFISH allowed us to subdivide IAF populations in a way not apparent in the human data.

**Figure 7:**
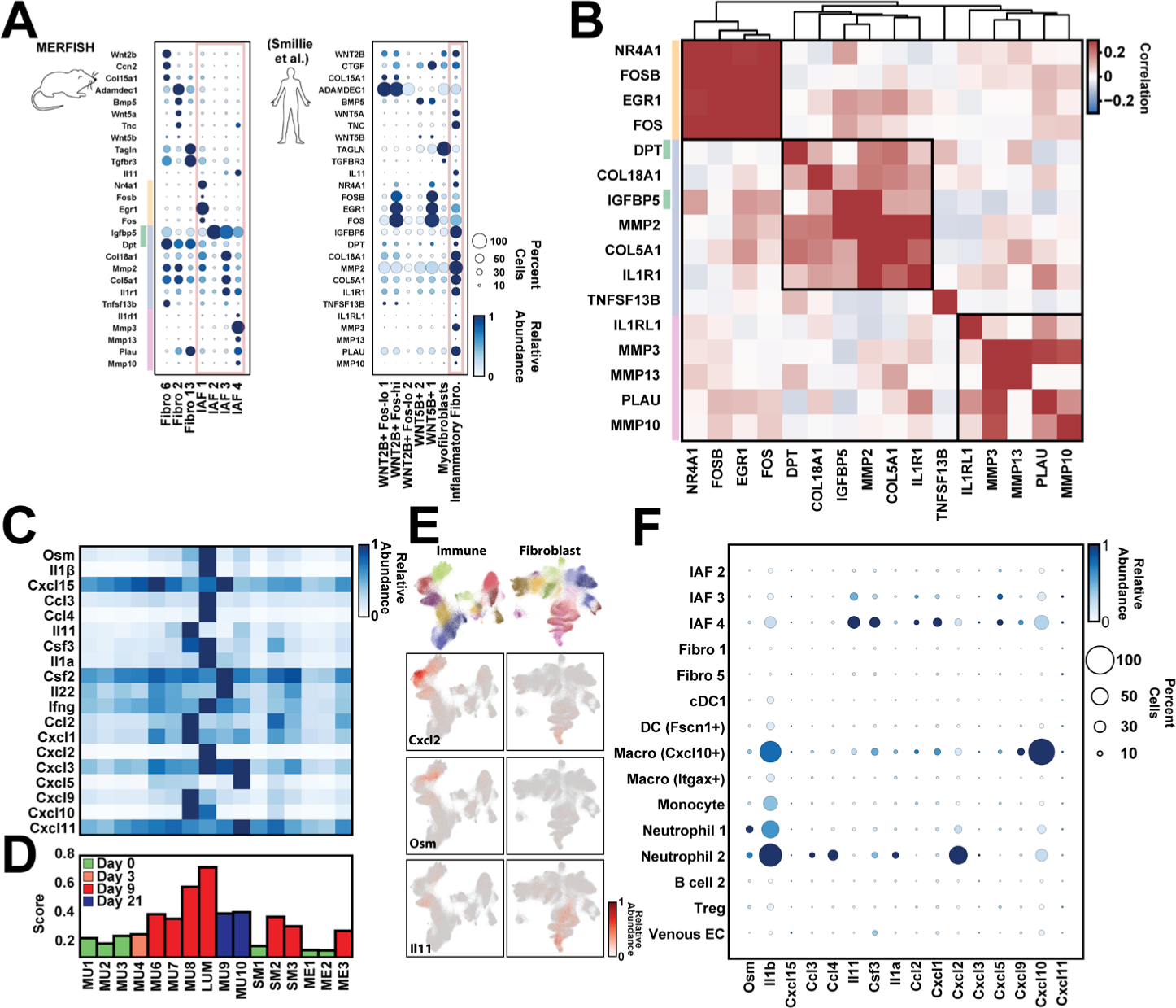
Signatures of murine IAF populations and late-stage inflammatory neighborhoods in human ulcerative colitis. (A) Average gene expression of key fibroblast markers in mouse fibroblast populations (left) or of their human homologs in published populations observed in UC biopsies (right). The color of the circle represents the average expression normalized to the maximum observed across all populations while the size of the circle represents the fraction of cells with at least one RNA copy. The red box highlights the IAF seen in mouse or human. The yellow, green, blue, and red lines represent the top mouse markers for IAF1 (yellow), IAF2 (green), IAF3 (blue), and IAF4 (red). (B) Pairwise Pearson correlation coefficients for the expression of the listed genes within the published human IAF population sorted hierarchically (top tree) with boxes indicating the genes grouped by this analysis. Colored bars on the left indicate the IAF population in mouse that these human homologs would mark as in (A). (C) The average expression of the mouse homologs of a human anti-TNF treatment- resistant UC biomarker panel in each of the listed mouse neighborhoods. Expression is normalized to the maximum observed in any neighborhood. (D) The biomarker score for each mouse neighborhood, with neighborhoods colored based on the disease stage at which they are most prominent. The biomarker score is defined as the average of the normalized expression for the markers listed in (C) in each neighborhood. (E) The mouse immune (left) and fibroblast (right) UMAPs colored either by the major cell label (top) or by the expression of the listed genes (bottom rows). (F) The average expression of each of the mouse homologs of human UC treatment resistant biomarkers in each of the major cell populations observed in MU8 and LUM as in (A).

If the human IAFs contained similar subpopulations, we reasoned that the markers of distinct mouse IAFs should covary within the human IAF cells. To explore this possibility, we computed the cross-correlation of these IAF-subset marker genes within human IAF cells, and, indeed, we found these markers organized into three covarying groups (**Figure 7B**), consistent with the existence of human IAF subpopulations. Remarkably, the groups of covarying genes in the human data agreed well with the mouse markers: with 4 of 4 IAF1 markers, 5 of 5 IAF4 markers, and 6 of 7 of the shared IAF2 and IAF3 markers comprising the covarying groups. The only proposed IAF subset marker that did not group coherently was *TNFSF13B* (BAFF), which was expressed at very low levels in human IAFs and correlated poorly with all other markers. Thus, this analysis suggests that at least some of the IAF subsets we observe in mouse may have human analogs in UC.

Building on this support for IAF subdivisions in humans and the role that IAFs play in mouse inflammatory neighborhoods, we next asked if there was evidence that similar neighborhoods might be present in human UC. To address this question, we leveraged a UC-disease-severity biomarker panel^72^, which predicts resistance to monoclonal anti-TNF treatment. We examined the expression level of the mouse homologs to panel members in each of our neighborhoods and observed that most biomarkers were strongly enriched in two neighborhoods—MU8 or LUM—with some enrichment in all neighborhoods seen at Day 9 or Day 21 (**Figure 7C-7D**). We noted similar results for a recently expanded UC biomarker panel^84^ (**Figure S7**). Interestingly, no single population of cells in MU8 or LUM produced all biomarkers, with marker subsets produced by a variety of immune populations (Macrophage [Cxcl10+], Monocytes, Neutrophil 1, and Neutrophil 2) or IAF3 and IAF4 (**Figure 7E-7F**). Thus, MU8 and LUM appear to be the predominant source of the mouse homologs of the human UC biomarkers, suggesting that analogous neighborhoods exist in human UC patients and that their presence correlates with anti-TNF treatment resistance. Importantly, this assertion agrees with the assignment of MU8 and LUM to ulcers, as the presence of ulcerated tissue features prominently in histological scores^85^ that predict treatment non-responsiveness^16^. These observations highlight one of the promises of spatial transcriptomic methods: the ability to merge the historically distinct spatial information provided by histology with the molecular and cellular information often captured with molecular biomarkers.

## DISCUSSION

IBD involves the contributions of multiple cell types present in healthy tissues and induced or recruited in the context of inflammation and tissue repair, all intricately organized into the structures seen in healthy, inflamed, or repairing tissues. To provide a measure of the cellular and spatial remodeling that occurs in IBD, we leveraged MERFISH to profile the expression of 940 genes in 1.35 million cells imaged during the onset of and recovery from DSS-induced colitis. The atlas that these measurements produced, which provides coverage of multiple pathways important for gut homeostasis and inflammatory response, opens multiple broad windows into the response of the gut to experimental colitis. Thus, we anticipate that this atlas could be further mined to explore a variety of questions and propose hypotheses for cell populations beyond those we considered here.

### Fibroblast diversity in the healthy and inflamed gut

Our MERFISH measurements defined cell populations within the expected cell classes in the healthy and inflamed gut, with expression profiles and divisions consistent with and extending prior work. Of these different cells, we focused, in part, on fibroblasts, where questions remain on their heterogeneity and location. By combining our measurements of gene expression and spatial location with the single-cell profiles provided by previous studies, we conclude that there is a consensus on the major fibroblast groups in the healthy colon. Three major groups are found within the lamina propria, one in the sub- mucosa, and one in between the inner and outer muscle layers. In addition, our measurements reveal gene expression variations within cells of many of these populations, with some variation cued to tissue location, consistent with further transcriptional and perhaps functional sub-divisions within these groups.

In addition, our data now suggest that this healthy fibroblast diversity is reflected in a similar diversity of inflammation-activated or polarized fibroblast states that emerge in the context of colitis from distinct healthy fibroblast groups. By leveraging a combination of shared gene expression, location, and targeted lineage tracing, we propose the following origins. IAF1 represents an early response state and arises from multiple healthy LP fibroblasts, which we do not distinguish either due to a modest number of cells, a similar transcriptional response, or the lack of critical genes in our MERFISH panel; IAF2 and IAF3 represent a polarized form of the crypt base fibroblast population, with IAF2 and IAF3 representing earlier and later stages in this polarization; IAF4 represents the polarized form of top-crypt fibroblasts; and Fibro 4 is a polarized form of the submucosa fibroblast group Fibro 7. We did not label Fibro 4 as an IAF, despite our proposal that it represents a polarized form of the submucosal fibroblast population, because a small number of these cells were found in healthy tissues near lymphoid follicles. The shared similarities in gene expression between these submucosal cells near follicles and those polarized during inflammation might reflect a similar function in enhanced immune recruitment.

One important consequence of distinct IAF populations is the ability to execute distinct inflammatory functions in different tissue regions. This point is exemplified by the IAF diversity within the MU8 ulcers, where at least three different IAF populations are found and are differentially distributed along the axis between the lumen and submucosa. The enhanced expression of ECM remodeling factors (e.g., *Mmp3*, *Mmp10*, and *Mmp13*) in IAF4 or collagens (e.g., *Col18a1*) in IAF2/3 suggests a functional partitioning between these IAF subsets with respect to ECM remodeling. Similarly, the enhanced expression of inflammatory cytokines (e.g., *Il1β* and *Csf3*), chemokines that recruit granulocytes (e.g., *Cxcl1* and *Cxcl2*) or T cells and DC (e.g., *Cxcl10*), and stromal activation factors (e.g., *Il11*) by IAF4 highlight the differential role this population plays in immune recruitment and activation as well as fibroblast polarization. The gene expression gradients observed within IAF4 further underscores the spatial partitioning of fibroblast functions and raises the possibility that further transcriptional and functional divisions may remain to be described. Activated fibroblast states have now been defined in a variety of contexts, including cancer, rheumatoid arthritis, and autoimmune disease^86–88^. As these states share similar gene expression signatures, IAFs may have similar roles across these disparate contexts^86–88^.

The increased IAF diversity we now describe raises a series of interesting questions. While our data suggest that healthy fibroblast groups differentially polarize to distinct IAFs, it is unclear if this polarization is always one-to-one or if additional polarized subsets might exist within these groups. For example, not all fibroblast groups in healthy tissue have a clear association with IAF populations in our data (e.g., Fibro 15 and Fibro 13), raising the possibility that some groups are lost in disease or polarize to similar activated states. Similarly, while we capture potential intermediate states in the fibroblast activation process (e.g., IAF1 and perhaps IAF2), the full fate and trajectory of healthy fibroblasts during inflammation and in the potential depolarization process during repair remains unclear. In addition, the molecular signals that drive fibroblast activation, as well as the relative contribution of host- or microbially derived factors, remain incompletely described. Fortunately, multiple aspects of our atlas, including a better definition of fibroblast and IAF markers, locations, and potential cell-cell communication channels, could prove useful in further dissection of these open questions. In parallel, by leveraging the markers of mouse IAFs, we identified statistical signatures of IAF subdivisions within human IL-11*+* IAFs from UC biopsies, raising the possibility that fibroblast activation is conserved in humans and these subdivisions may play similar functional roles to those we defined in mice. Importantly, our studies suggest that ulcerated tissues would be the most informative tissue structure for further study of these potential human subdivisions.

### Staged inflammatory tissue remodeling

Gut inflammation has long been associated with a series of defined morphological changes in gut structures. Our measurements now provide the cellular and molecular underpinnings of these structural changes, revealing that the spatial remodeling of the gut during inflammation is defined by the emergence of a series of spatially ‘patchy’ cellular neighborhoods with sharp spatial boundaries. Our analysis further suggests these neighborhoods emerge in an ordered temporal progression, with distinct responses in the mucosa, submucosa, and muscularis externa. In this light, the increased slice-to-slice variability observed in Day 9 tissue may be a result of the differential progression of different mucosal regions along this tissue inflammation axis. By contrast, a similar linear progression cannot be suggested for the Day 21 neighborhoods, as repair starts from the highly variable tissue structure observed at Day 9. Importantly, this staged progression, which we proposed solely based on molecular and cellular measures of the tissue, agrees nicely with the progression identified throughout purely histological measures of IBD.

The patchy-like distribution of cellular neighborhoods in combination with their sharp spatial boundaries suggests that there are positive reinforcement mechanisms that drive neighborhood composition. For example, cells within a neighborhood might secrete chemokines that recruit the same defining cell populations to the surrounding tissue, reinforcing neighborhood identity. Alternatively, signals generated within the neighborhood could further polarize tissue resident cells, like fibroblasts, in a way that reinforces neighborhood identity. One such mechanism would be the polarization of fibroblasts by *Il11* produced by IAFs, as suggested by others^80–82, 89^ and our receptor-ligand analysis. Nonetheless, the mechanisms that drive neighborhood formation and progression remain to be defined. On this front, the receptor-ligand interactions we suggest could provide a useful resource for directing further studies.

Finally, we propose that mouse neighborhoods may have analogs in human IBD. We show that the mouse homologs of human biomarkers for severe IBD—as judged by treatment non-responsiveness—are strongly enriched in mouse ulcerated neighborhoods, suggesting that, indeed, many features of the mouse ulcer are conserved in human. Supporting this assertion, recent work^16^ has revealed UC ulcers as a major source of these human biomarkers and through gene-module analysis proposed that human ulcers contain neutrophils, IL-1β+ mononuclear phagocytes, IL-11+ activated fibroblasts, and increased vascularization—features we see in MU8. Interestingly, in CD a similar collection of cells, comprising plasma cells, inflammatory mononuclear phagocytes, activated T cells, and activated fibroblasts has been identified and associated with treatment non-responsiveness^30^, suggesting some conservation in neighborhood composition in severe UC and CD. Here we have leveraged MERFISH to map the inflammatory response in a mouse model, and, given the insights we provided in the mouse response, an understanding of human IBD might benefit from similar measurements. More broadly, the ability to chart the molecular, cellular, and spatial organization and reorganization throughout disease by the approaches we leveraged here could offer a broad range of insights in a multitude of inflammatory contexts.

## Acknowledgements

We thank members of the Moffitt and Nowarski laboratories as well as José Ordovás- Montañés for helpful discussion and critical reading of the manuscript. We thank Dr. Charles G. Jennings and Lai Ding for their support. This work was supported by grants made available to R.N. by the Kenneth Rainin Foundation, NIH (R35GM133800), and Crohn’s and Colitis Foundation; to J.R.M. from the Chan Zuckerberg Initiative, the Helmsley Charitable Trust, and the NIH (R01GM143277 and P30DK034854); to K.N.S. by the National Health and Medical Research Council (NHMRC) Australia through the CJ Martin Overseas Biomedical Early Career Fellowship (GNT1143655); and to A.M. by the Department of Defense (DoD) through the National Defense Science & Engineering Graduate (NDSEG) Fellowship Program. Measurements in this manuscript were supported by the Neurotechnology Studio at Brigham and Women’s Hospital.

## Author contributions

Conceptualization, P.C., K.N.S., J.R.M., and R.N.; Methodology, P.C., K.N.S., A.M., R.J.X., D.M., E.Y., J.M.R., K.T., Y.K., L.B., G.W., L.S., J.R.M., and R.N.; Software, P.C., R.J.X., L.G., T.L, and J.R.M.; Investigation, P.C. and K.N.S.; Formal Analysis, P.C. and K.N.S.; Data Curation, P.C. and K.N.S.; Resources, N.O., F.J.Q., A.C.A., V.K.K., J.R.M., and R.N.; Writing – Original Draft, P.C., K.N.S., J.R.M., and R.N.; Writing – Review & Editing, P.C., K.N.S., A.M., R.J.X., D.M., E.Y., J.M.R., K.T., Y.K., L.B., L.G., T.L., N.O., G.W., L.S., F.J.Q., A.C.A., V.K.K., J.R.M, and R.N.; Visualization, P.C., K.N.S., J.R.M., and R.N.; Supervision, J.R.M. and R.N.; Funding Acquisitions, J.R.M. and R.N.

## Supplementary Figures and Legends

**Figure S1:**
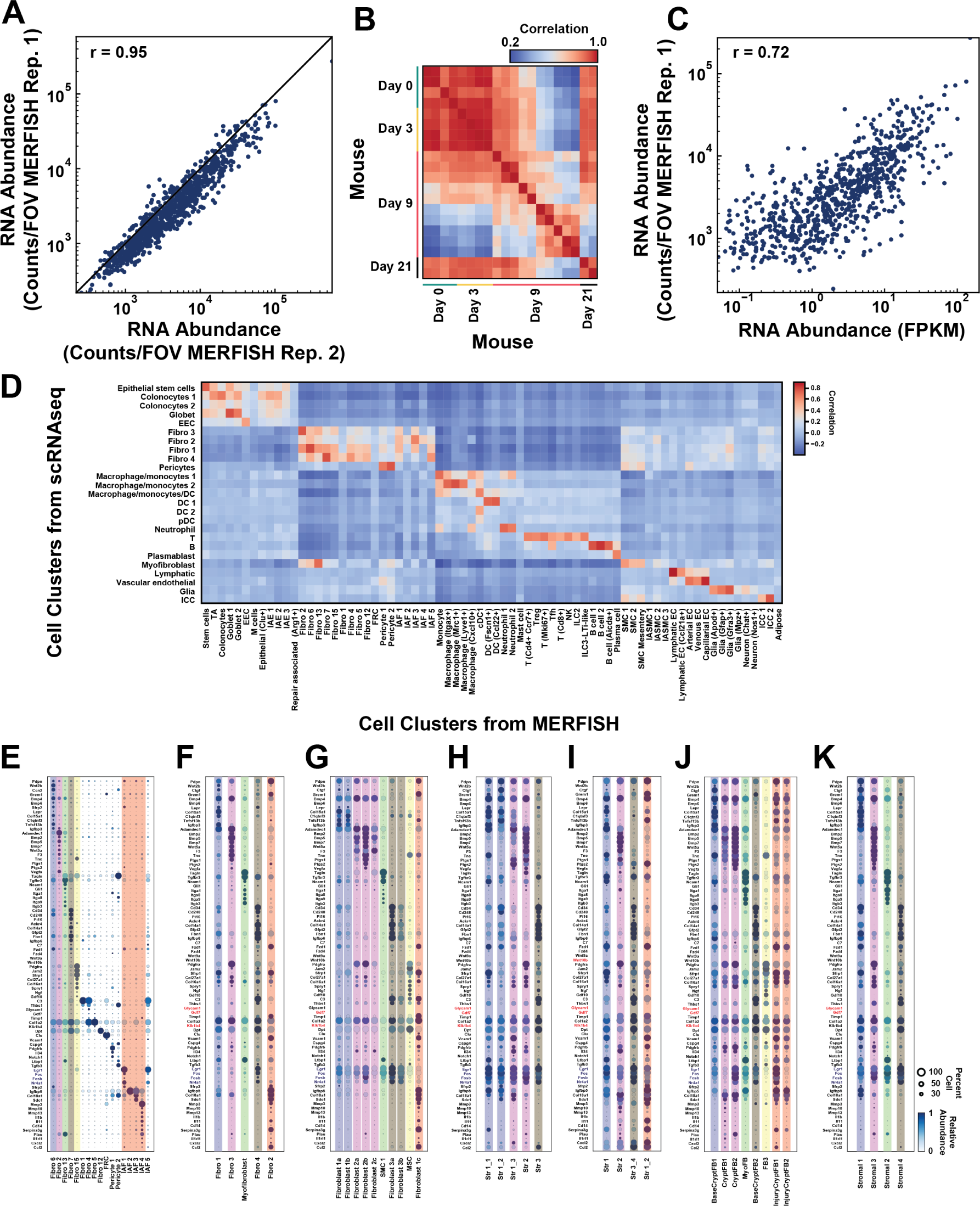
Reproducibility and cross-validation of MERFISH measurements. (A) Scatter plot of counts per cell for each RNA averaged across all fields-of-view (FOV) for one MERFISH measurement in a Day 0 mouse versus those for a replicate MERFISH measurement in a different Day 0 mouse. The Pearson correlation coefficient (r) between the logarithm of the average expression values is 0.95. (B) Heat map of the pairwise Pearson correlation coefficients calculated as in (A) for all MERFISH measurements (each measurement often contains multiple slices), indicating strong reproducibility between measurements of mice from different disease stages. (C) Scatter plot of the average counts per cell for each RNA measured with MERFISH from a representative Day 0 replicate versus the average expression determined from bulk RNA sequencing of tissue harvested from Day 0 mice. The Pearson correlation coefficient between the logarithm of the average expression values is 0.72. (D) Heatmap of the pairwise Pearson correlation coefficients determined between the expression profiles measured for cell clusters identified in MERFISH and those previously determined via scRNAseq^39^. Expression was determined by averaging the z-score of the logarithmic expression for all cells within a given cluster. (E) Dot plot representation of the expression of key genes for fibroblast populations determined with MERFISH. Color indicates the relative expression of each gene across the listed cell populations while the circle size represents the fraction of cells with at least RNA copy. The colored bars represent corresponding fibroblast populations observed in published data as judged by the co-expression of key marker genes. (F-K) As in (E) but for all fibroblast populations identified via scRNAseq of the mouse colon in healthy and DSS-induced colitis from Ho et. al.^39^ (F), Jasso et. al.^18^ (G), Kinchen et. al. (healthy)^9^ (H), Kinchen et. al. (DSS-treated)^9^ (I), Xie et. al.^40^ (J), and Brügger et. al.^15^ (K).

**Figure S2.**
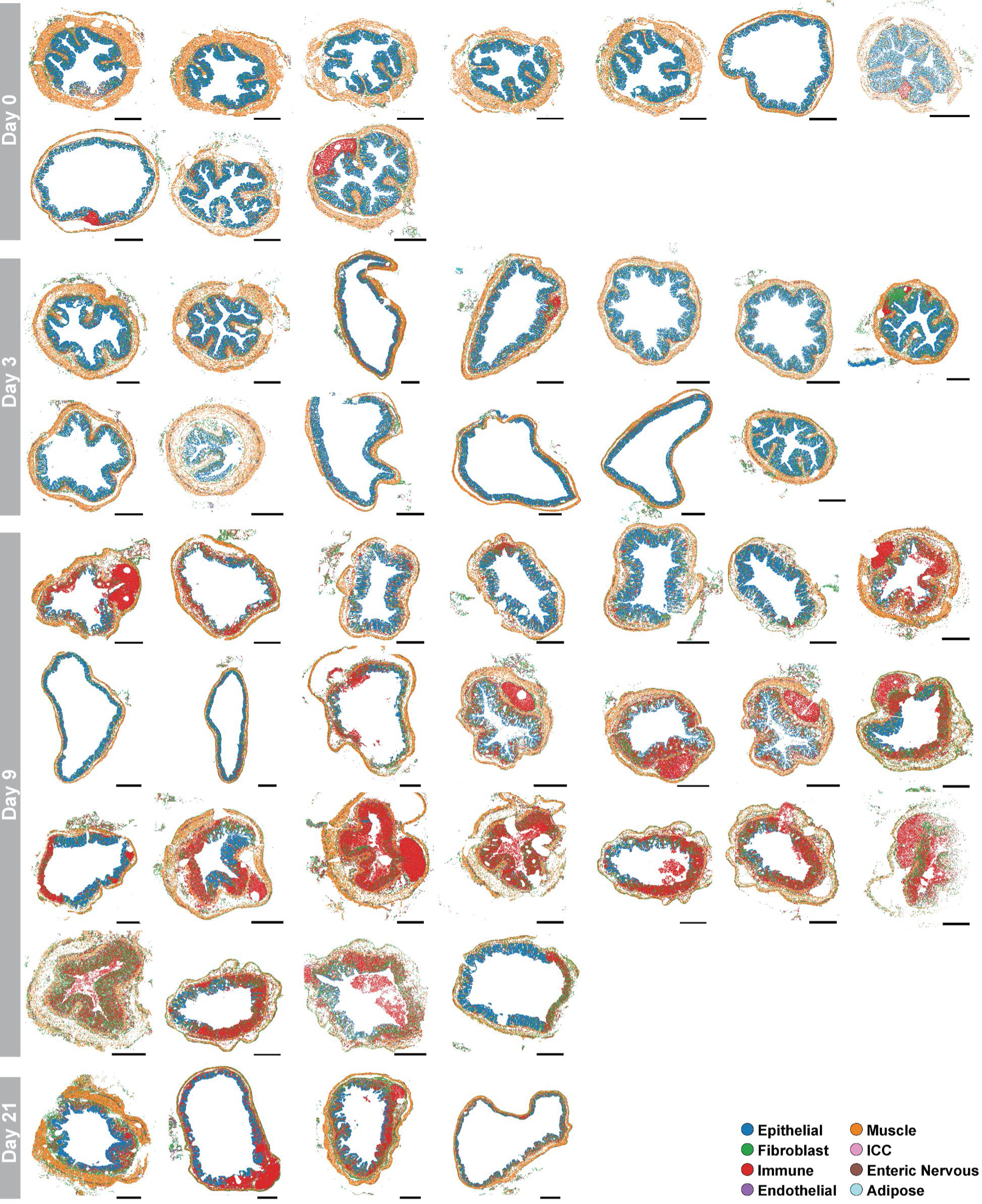
The spatial distribution of cell classes in all measured slices. Each dot represents the center of a cell, and the color indicates the cell class assignment. Slices are grouped in rows by the disease stage from which they were collected. Scale bars: 500 µm.

**Figure S3.**
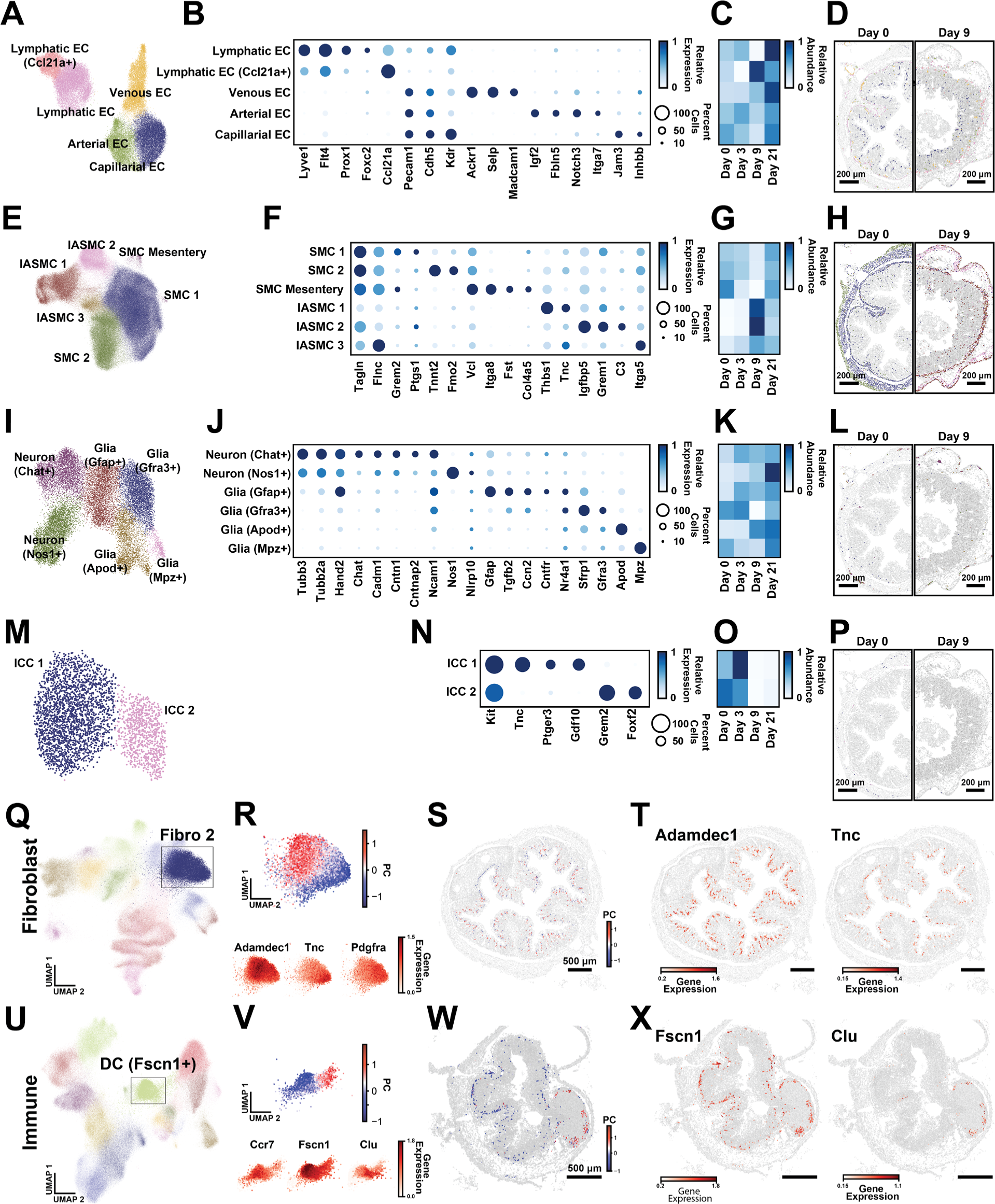
Gene expression and spatial distributions for sub-populations within major cell classes identified with MERFISH. (A) UMAP representation of the endothelial cells (EC), colored by identified clusters. (B) Dot plot representation of key marker genes for the populations of EC. The color indicates the relative expression of each gene across all listed populations, and the size of the marker indicates the fraction of cells that express at least one copy of that RNA. (C) The fractional abundance of each of the identified EC populations across disease stage, normalized to the maximum abundance observed at all disease stages. (D) The spatial distribution of EC in representative slices taken from Day 0 and Day 9. Markers represent the center of cells. Gray represents all cells, and all EC are colored corresponding to their cluster identities as shown in the UMAP in (A). Scale bars: 200 µm. (E-H) As in (A-D) but for smooth muscle cells (SMC). (I-L) As in (A-D) but for cells of the enteric nervous system. (M-P) As in (A-D) but for interstitial cells of Cajal (ICC). (Q) UMAP representation of fibroblasts cells highlighting the cells assigned to Fibro 2. (R) Zoom in on the boxed region of the UMAP in (Q) with cells colored by the amplitude of a principal component (PC) associated with a PC analysis of gene expression within cells assigned to Fibro 2 (top) and colored by the expression level of three genes that contribute to the loading associated with this PC (bottom). (S) Spatial distribution of Fibro 2 cells within a representative slice. All cells are colored gray, and all cells assigned to Fibro 2 are colored by the amplitude of the PC as seen in (B). Scale bar: 500 µm. (T) Spatial distribution of Fibro 2 cells colored by the expression of two genes that define the loading associated with the highlighted PC in (R). Scale bar as in (S). (U-X) As in (R-T) but for DC (Fscn1+).

**Figure S4.**
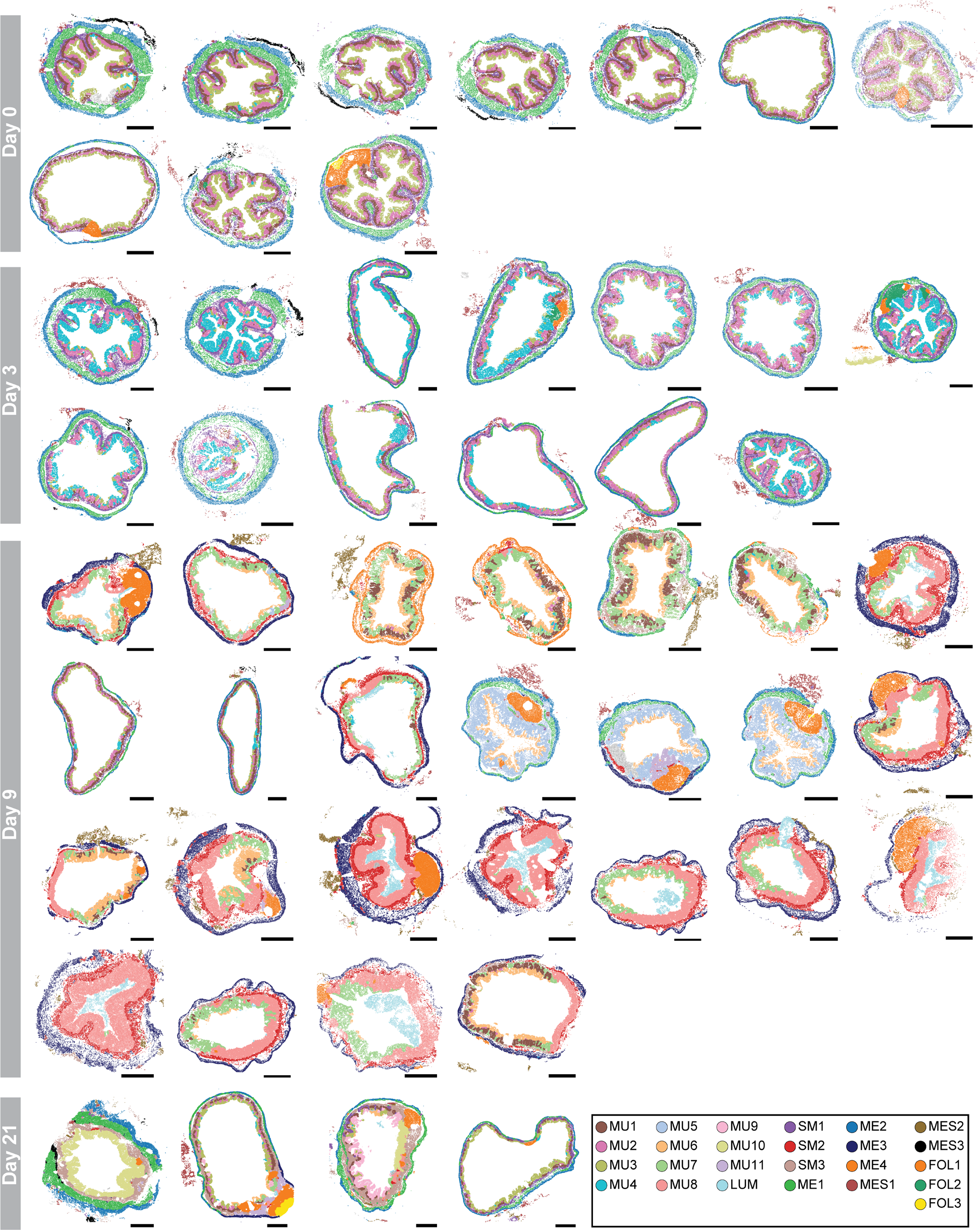
The spatial distribution of cellular neighborhoods in all slices. Each dot represents the center of individual cells and color indicates the neighborhood assignment. The rare gray cells represent the 0.5% of cells not assigned to one of our neighborhoods. Slices are grouped in rows by the disease stage from which they were collected. Scale bars: 500 µm.

**Figure S5.**
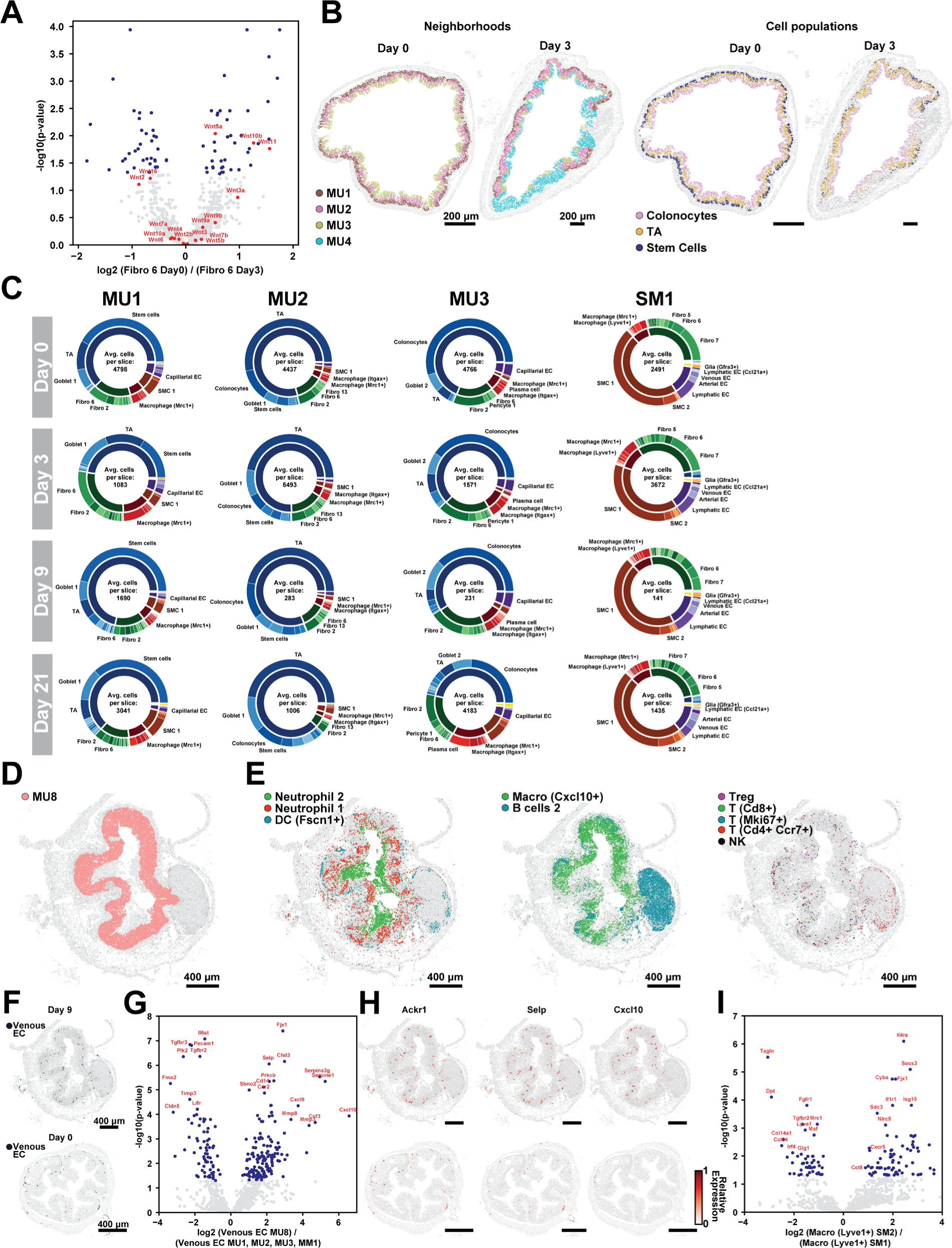
Additional properties of cellular neighborhoods. (A) Volcano plot of the differential expression of genes in Fibro 6 cells seen at Day 3 versus that seen at Day 0. Genes highlighted in blue pass a false discovery threshold of 5% while named genes marked in red represent all Wnt-family members. (B) The spatial distribution of cells in a representative Day 0 and Day 3 slice with cells colored by the cellular neighborhood to which they were assigned (left) or the cell population to which they were assigned (right). Scale bars: 200 µm. (C) Pie charts representing the abundance of cell types in MU1, MU2, MU3, and SM1 for only the cells associated with Day 0, Day 3, Day 9, or Day 21. Pie charts are displayed as described in Figure 3. (D, E) The spatial distribution of cells in a representative Day 9 slice colored by cells assigned to MU8 (left) or major clusters found in MU8 (right). Scale bars: 400 µm. (F) The spatial distribution of cells in a representative Day 9 (top) or Day 0 (bottom) slice with Venous EC cells in blue. Scale bars: 400 µm. (G) Volcano plot of the expression of all genes within Venous EC in MU8 or that in MU1, MU2, MU3, or MM1 as in (A). (H) The spatial distribution of all cells in the representative slices shown in (F). Only Venous EC cells are colored, and they are colored based on the relative expression of the listed genes. (I) Volcano plot of the differential expression of genes in Lyve1+ macrophages see in SM2 versus those seen in SM1.

**Figure S6.**
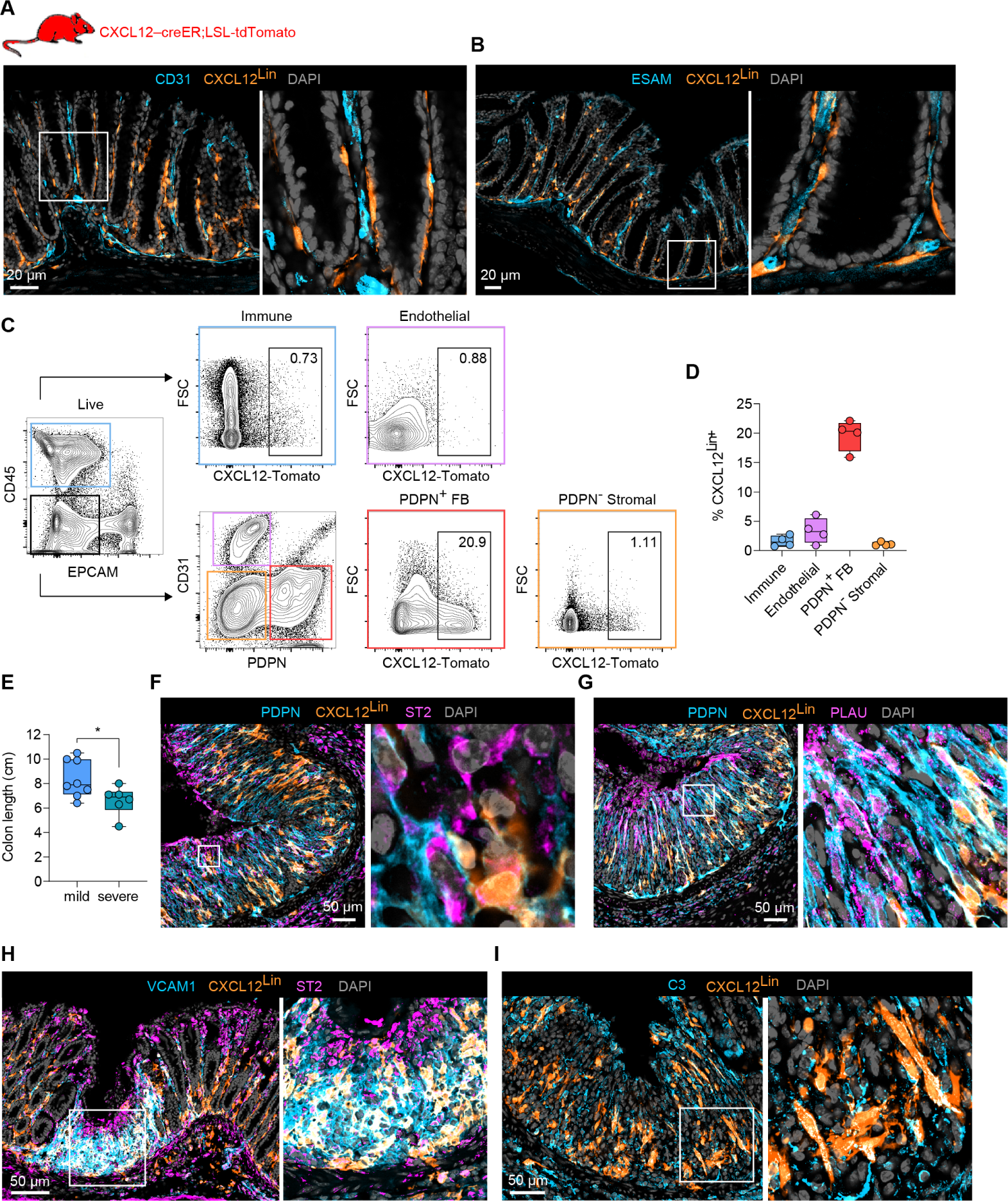
Lineage tracing reveals distinct origins of inflammation-associated fibroblasts. (A, B) Representative immunofluorescence images of distal colon from healthy CXCL12^Lin^ mice (CXCL12-creER; LSL-tdTomato; *n*=5) for CD31 (A) or ESAM (B). Scale bars: 20 µm. (C, D) Flow cytometry analysis of colonic CXCL12-Tomato expression in immune (CD45+), endothelial (CD31+), PDPN+ fibroblasts (PDPN+ +CD45-CD31-EpCAM-) and PDPN- stromal (PDPN-CD45-CD31-EpCAM-) cells at steady-state (*n*=6). (E) Colon length measured for the CXCL12^Lin^ mice sorted into groups that had mild or severe degrees of weight loss as measured at Day 9 (*n*=14). The center line represents mean, the width of the bars represents standard deviation, and the whiskers extend to include 95% of the data. Individual markers represent distinct mice. *p<0.05. (F-I) Representative immunofluorescence images of inflammatory regions in distal colon from mice, harvested on Day 9 (n=6). The degree of overlap between the CXCL12^Lin^ marker and other markers of IAF populations, ST2 (F), PLAU (G), VCAM1 (H) and C3 (I), are highlighted. Scale bars: 50 µm.

**Figure S7.**
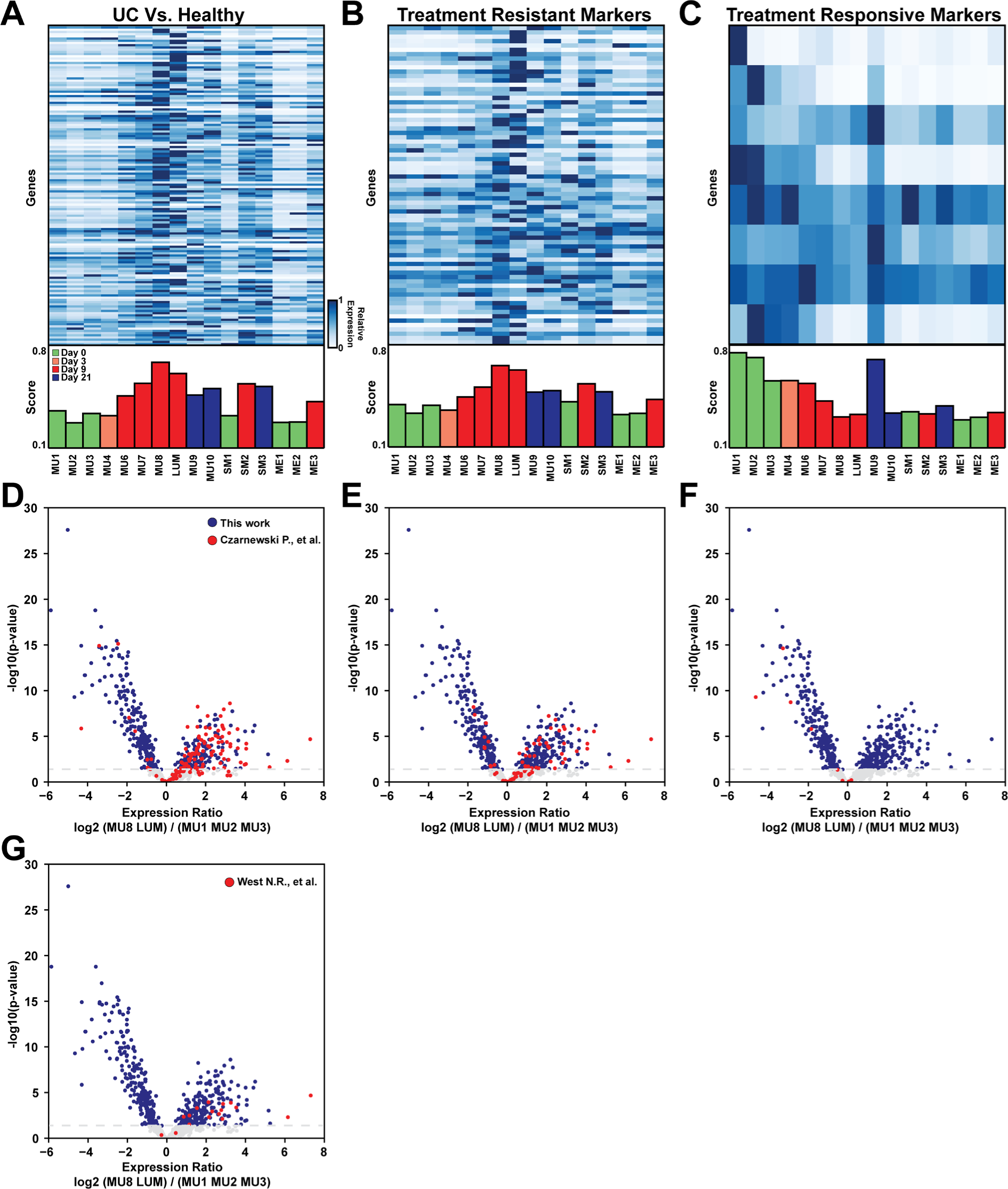
Human biomarker panel expression in mouse neighborhoods supports the presence of similar neighborhoods in human ulcerative colitis. (A-C) The average expression of the mouse homologs of the human biomarkers reported by Czarnewski et. al.^84^ to distinguish UC from healthy samples (A), anti-TNF resistant UC samples from anti-TNF sensitive samples (B), anti-TNF sensitive samples from anti-TNF resistant samples (C), within the listed mouse neighborhoods (top) or the average biomarker score (as in Figure 7) for these neighborhoods (bottom). (D-G) Volcano plot of the differential expression of mouse genes observed in the MU8 and LUM neighborhoods relative to that observed in the healthy neighborhoods, MU1, MU2, and MU3. Blue marks all genes that pass a false-discovery-rate threshold of 5% (dashed gray line), and red marks the mouse homologs of the human biomarkers reported by Czarnewski et. al.^84^ to distinguish UC from healthy samples (D), anti-TNF resistant UC samples from anti-TNF sensitive samples (E), anti-TNF sensitive samples from anti-TNF resistant samples (F), or from the anti-TNF-resistant UC biomarkers reported by West et al.^72^.

